# Client distribution between Chlamydomonas FDX1 and FDX2 in carbon, nitrogen and sulfur assimilation

**DOI:** 10.64898/2026.01.14.699543

**Authors:** Stefan Schmollinger, George Kusi-Appiah, David Villegas, Sarah Stainbrook, Thomas V. O’Halloran, Daniela Strenkert

**Affiliations:** Department of Energy Plant Research Laboratory, Michigan State University, East Lansing, MI, 48824; Department of Biochemistry and Molecular Biology, Michigan State University, East Lansing, MI, 48824; Department of Microbiology, Genetics and Immunology, Michigan State University, East Lansing, MI, 48824; Department of Chemistry, Michigan State University, East Lansing, MI, 48824; Department of Plant Biology, Michigan State University, East Lansing, MI, 48824

## Abstract

Plant-type ferredoxins (Fd) comprise small, soluble protein families that distribute electrons from photosystem I to various client proteins within the chloroplast stroma. In *Chlamydomonas reinhardtii*, the major, constitutively expressed FDX1/PetF supports Fd-NADP^+^ reductase (FNR) in NADPH production. The highly similar FDX2 is present only when its preferred nitrogen (N) source ammonium is absent, supplying Fd-dependent nitrite reductase (NiR) for nitrate/nitrite assimilation. Surprisingly, despite accumulating to ∼10% of FDX1 abundance and preferential interaction with NiR, *fdx2* mutants are asymptomatic when grown on nitrate, requiring to additionally deplete FDX1 for growth to be halted. A *fdx1* knockout itself appears lethal, severe *fdx1* knockdowns have reduced growth rates both in phototrophic and photoheterotrophic conditions, independent of the N source. Transcriptome analyses of *fdx1* mutants revealed expression patterns similar to sulfur (S) deficient algae, and *fdx1* strains have a reduced total cellular S content. S assimilation requires Fd-dependent sulfite reductase (SiR) activity, an enzyme distantly related to FDX2 client NiR. Expression defects are partially alleviated; growth and S content are less impacted with FDX2 expression. Our mutant analysis shows the two major Fds in Chlamydomonas focus on a specific subset of Fd-dependent metabolism, mostly supplying Fd-dependent enzymes involved in macronutrient assimilation (C/N/S).

## INTRODUCTION

Ferredoxins (Fd) are conserved proteins in bacteria, eukaryotes and archaea that provide electrons to a wide range of metabolic reactions, often involving multiple Fd isoforms in the same organism (1–3). Since protein electron carriers evolve, domains can be duplicated or combined with others, and each individual structure can adapt via amino acid changes to either adjust their redox potential or better distinguish interaction interfaces, improving their utility for different client proteins and reactions (4). Fds are ancient Fe-S cluster proteins, allowing the delivery of one electron at a time (5, 6). They are classified based on the number and type of Fe-S cluster they house ([2Fe-2S], [3Fe-4S] or [4Fe-4S]), and more closely based on the distribution of the cysteines that are involved in the binding of these Fe-S clusters to the Fds (7, 8). Ancestral Fds were likely [4Fe-4S] proteins, these clusters can form naturally in the low oxygen environments where early life evolved, that are more sensitive to oxygen compared to [2Fe-2S]-containing isoforms (7, 9, 10). This preference likely contributed to selection for [2Fe-2S] cluster usage in oxygenic environments after the great oxidation event and in oxygenic phototrophs (11).

Plant-type Fds are a subgroup of [2Fe-2S] Fds localized to eukaryotic plastids, distinguished by their small size (∼ 10 kDa), a high proportion of acidic amino acids, a characteristic cysteine spacing pattern (CX_4_CX_2_C) and a relatively negative redox potential (12–14). While cyanobacteria already employ a suite of Fds equipped with varying Fe-S cluster types, eukaryotic phototrophs almost exclusively use [2Fe-2S] clusters (15). During oxygenic photosynthesis, electrons that are extracted from water at PSII mainly follow linear electron flow (LEF), with stroma-facing PSI subunits ultimately transferring the electrons to Fds (16, 17). Most electrons are directed by Fd towards Fd–NADP+ oxidoreductase (FNR), reducing NADP+ to NADPH, which in turn is used mostly in the Calvin cycle (CBB) during the reduction of 1,3-bisphosphoglycerate to glyceraldehyde 3-phosphate (18, 19). In non-photosynthetic cells/tissues of plants, or during the night in algae, Fd reduction necessitates a reversal of the FNR reaction, transferring electrons from NADPH, which in turn can be generated in the oxidative pentose phosphate pathway or from other sources (20–22). In the eukaryotic algae *Chlamydomonas reinhardtii* (Chlamydomonas hereafter), a reference organism in the green lineage (Viridiplantae) to study eukaryotic photosynthesis (23), the highly abundant and conserved FDX1/PetF (FDX1 hereafter) is the likely Fd to deliver electrons to FNR (24). Fds also contribute to cyclic electron flow (CEF) around PSI, which transfers electrons back to the cyt *b*_6_*f* complex, directing electrons away from NADPH production and instead promoting the proton gradient across the thylakoid membrane, balancing photosynthetic ATP vs NADPH production (24–27).

The multiple Fd isoforms evolved in cyanobacteria were to some degree retained and expanded on in eukaryotic algae and land plants after endosymbiosis (28). Sequence similarity between individual Fd isoforms varies widely (29), which may reflect evolutionary advantages from, both, some functional redundancy between closely related Fds, but also neo-functionalization and specialization of Fds with larger differences in protein sequence (30). This is reflected by the observation that a single, major Fd in chloroplasts appears to account for most of LEF, while other Fd isoforms vary widely in abundance and can be conditionally expressed only in specific environments (31, 32). Many metabolic processes require Fds for electron supply in the chloroplast, most equally essential for photosynthetic growth, including nitrogen and sulfur assimilation (33–35), fatty acid metabolism (36), pigment biosynthesis (37–39), the maintenance of chloroplast redox balance via the thioredoxin system (40) and, in eukaryotic algae, hydrogen production (41). The *Chlamydomonas reinhardtii* genome codes for a total of up to 12 chloroplast-targeted Fd isoforms including the major, photosynthetic Fd, FDX1 (42).

Nitrogen (N) is an essential element for all living organisms and is frequently bio-unavailable in the environment, limiting plant growth and photosynthetic productivity (43). Ammonium (NH_4_^+^), nitrate (NO_3_^−^) and nitrite (NO_2_^−^) are the most frequent inorganic N sources assimilated by photosynthetic eukaryotes, including in aquatic systems (44). Like other microalgae Chlamydomonas has a flexible metabolism with respect to N assimilation and can utilize a variety of different N sources, with ammonium being preferred and nitrate being the second most desired N source (45, 46). Nitrate assimilation is accomplished by two transport and two reduction steps: import of nitrate into the cytosol where its reduced to nitrite, which is imported into the chloroplast and further reduced to ammonium (46–51). Fd-dependent nitrite reductase (NiR) completes this last step of the reduction process, requiring the delivery of all 6 required electrons by the 1-electron donor Fd. NiR is an Fe enzyme utilizing an unusual, closely linked siroheme and [4Fe-4S] cluster cofactor to accommodate this reaction (52, 53). Following reduction, NH_4_^+^ is assimilated via the chloroplast glutamine synthetase / glutamate synthase (GS/GOGAT) cycle (45, 54), where ammonium is loaded by GS onto glutamate (Glu) to produce glutamine (Gln), before 2 molecules of Glu are regenerated at GOGAT from 2-oxoglutarate and Gln, assimilating one N in the process. Phototrophic organisms contain a chloroplast localized GOGAT enzyme that also utilizes electrons from Fds during NH_4_^+^ assimilation (55, 56). Unlike NiR, GOGAT is required for N assimilation independent of the N source and, while NiR is exclusively Fd-dependent, a second, alternative GOGAT isoform is expressed in Chlamydomonas chloroplasts that can utilize NADH instead of Fd for electron supply (57, 58). FDX2 likely serves as the predominant ferredoxin (Fd) responsible for delivering electrons to NiR for nitrite reduction, it is almost exclusively expressed when cells are cultivated with nitrate as their sole source of nitrogen (59, 60). FDX2 expression is also induced in gametes and gametogenesis is induced by N- and S-deficiency in Chlamydomonas (61, 62). Additionally, FDX2 transfers electrons more efficiently to NiR than FDX1 *in vitro* (59).

Sulfur (S) is equally essential for life and, in eukaryotic phototrophs, its assimilation takes place in the chloroplast, requiring electrons from Fds (63). S is mostly assimilated from inorganic sulfate (SO_4_^2-^), which is imported to the chloroplast by a series of transporters (64–68). In the chloroplast, SO_4_^2-^ is assimilated into adenosine-5’-phosphosulfate/adenylyl-sulfate (APS) by ATP sulfurylases (ATS), before it is sequentially reduced to sulfide (S^2-^) (69–72). Similar to NiR, the second reduction step from sulfite (SO_3_^2-^) to sulfide requires 6 electrons, catalyzed by sulfite reductases (SiR), again involving a linked siroheme and [4Fe-4S] cluster (52, 53). While none of Chlamydomonas three SiR enzymes have been studied in detail, two of them require electrons from Fds, while the third one, SIR3, is a component of a bacterial-type enzyme receiving electrons from NADPH instead. While all three proteins are expressed in Chlamydomonas, the two Fd-dependent SiRs are increasingly expressed in response to S deficiency, the bacterial-type SiR is not (72). SiR and NiR share distant sequence similarity, show structural commonalities and *in vitro* can reciprocally reduce either ones substrates, NO_2_^−^ or SO_3_^2-^ (73, 74), albeit their kinetics indicate that *in vivo* each enzyme may be selective for its respective substrate. The resulting S^2-^ moiety from the SiR reduction is then assimilated onto a carbon skeleton (O-acetylserine) to produce the amino acid cysteine, which can then be used to produce methionine, glutathione or Fe-S clusters (75–78). Fd-independently, APS can already be used directly to produce 3’-phosphoadenosine 5’-phosphosulfate (PAPS), which in turn is used as a substrate of sulfotransferases to produce various S-containing metabolites; while sulfite (SO_3_^2-^) can be utilized to produce important sulfur containing lipids (79, 80).

Predicting Fd function can be challenging from sequence or expression alone; while Fds group well into subgroups over evolutionary scales (15, 28), the functional specialization of these groups is not well understood, electron carrier and clients can co-evolve in individual species and their function may vary in different ecological contexts. Reduction potentials between individual, plant-type Fds are often similar and *in vitro* activities can be misleading, requiring additional cellular context to provide specificity, including expression profiles, localization and considerations of abundance/composition of the cellular Fd pool during catalysis (81). To add to the information gathered on Chlamydomonas Fds, we chose a genetic approach to explore the contributions of the two most similar Chlamydomonas Fds (FDX1 and FDX2) to chloroplast metabolism.

## RESULTS

### FDX1 and FDX2 are highly similar, but differ in their charge profile

The occurrence of small families of chloroplast-targeted, plant-type ferredoxin (Fd) isoforms is a conserved feature within the green lineage. While this feature was already present in cyanobacteria, eukaryotes further expanded on the available structures, presumably to more efficiently supply electrons to their specific set of client proteins. The annotation of the Chlamydomonas genome contains 12 chloroplast Fds (*FDX1-12*), experimentally the chloroplast localization was confirmed for five of them (FDX1, FDX2, FDX3, FDX5 and FDX6 (59, 82). Eukaryotic algae are a diverse group of organisms, occupying marine, freshwater and soil environments, as well as more specialized niches. We first explored which of the different Fd isoforms present in Chlamydomonas are conserved amongst algae and focused our analysis primary endosymbionts. The N-terminus of the mature FDX1 protein has been determined experimentally (83), which we used to inform predictions from PredAlgo (84) and TargetP (85) to determine those of the other Chlamydomonas Fds (Data S1). Chloroplast targeting was predicted for 10 of the 12 Fds, all except for FDX10 and FDX12, both of which were also most divergent in sequence, also involving the cysteine spacing required for [2Fe-2S] cluster binding. A full alignment of all 12 Chlamydomonas Fds is provided in Figure S1A, while the region most similar to FDX1 for the 10 most similar Fds is depicted in Figure 1A.

**Figure 1.**
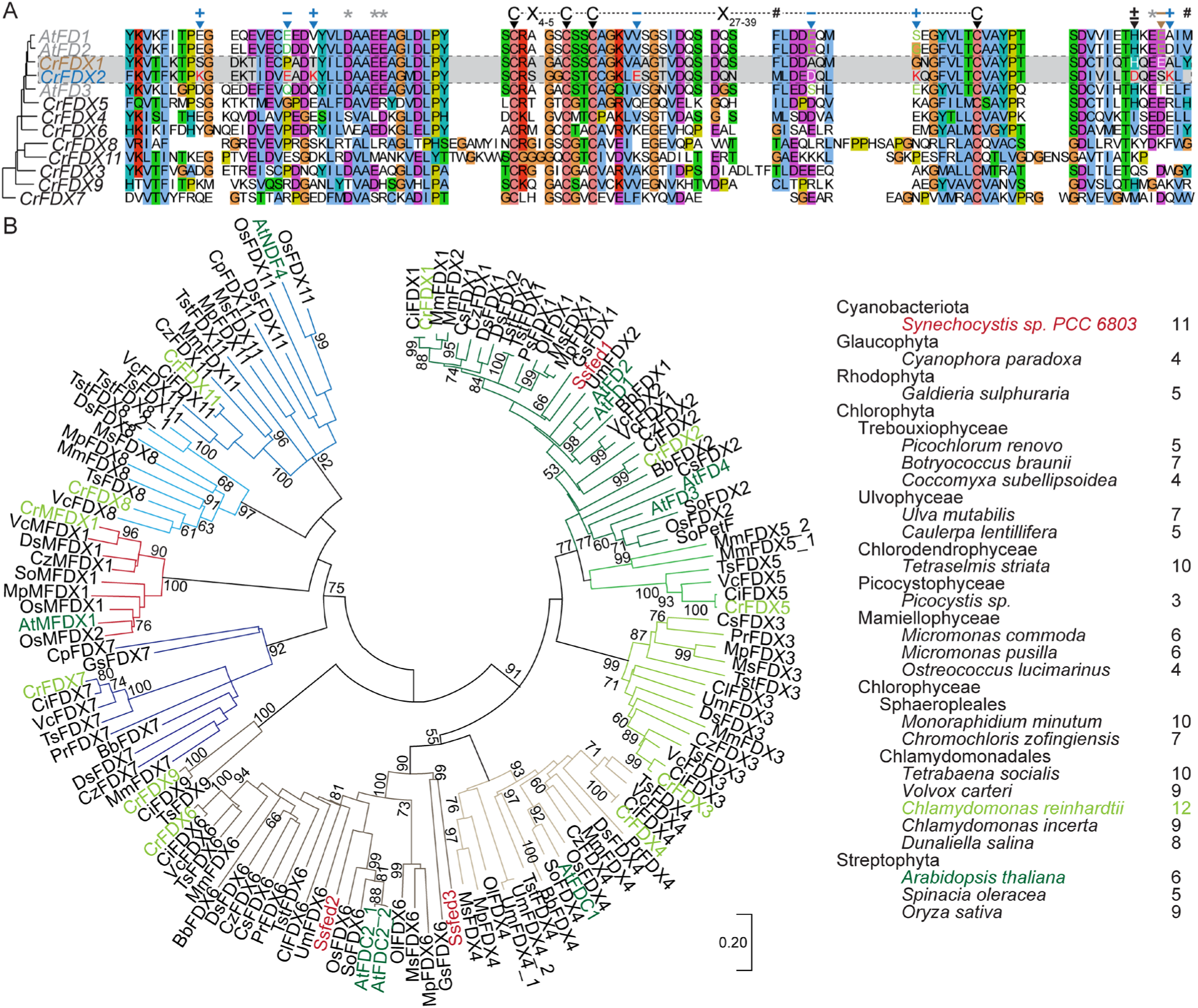
FDX1 and FDX2 are highly similar. (A) Multiple sequence alignment of the core Fd fold between root (FD3) and leaf (FD1/2) Fds from *Arabidopsis thaliana* and predicted chloroplast localized Fds from Chlamydomonas. Chloroplast transit peptides were removed prior to analysis, all sequences before and after processing are available in SData 1. Chlamydomonas FDX1 and 2 are highlighted with a grey background, changes from an uncharged to a charged amino acid in FDX1/2 are highlighted. Cysteines involved in 2Fe-2S cluster binding are indicated above. Stars indicate crucial positions for binding of FDX1 with FNR, hashes indicate mutants analyzed in (M. Boehm *et al.* (2015)). (B) Evolutionary conservation of Chlamydo-monas Fds among other eukaryotic algae. Ferredoxin families from various algae genomes were retrieved from Phytozome 14 (https://phytozome-next.jgi.doe.gov/) and Pytcocosm (https://phycocosm.jgi.doe.gov/phycocosm/home), the number of identified sequences are shown next to the organism in the legend (right panel). Individual alignments for each Chlamydomonas Fd were made to identify homologs in other algae, summarized in Figure S1. The subset of Fds from other algae, plants and Synechocystis that associated with Chlamydomonas proteins were combined and aligned using MAFFT. The complete alignment was used to construct a phylogenetic tree using the minimum evolution method in MEGA. The percentage of replicate trees in which the associated taxa clustered together in the bootstrap test (1,000 replicates) are shown next to the branching point. The tree is drawn to scale, with branch lengths in the same units as those of the evolutionary distances used to infer the phylogenetic tree.

All 10 of the analyzed Chlamydomonas Fds contain the necessary cysteines required for [2Fe-2S] cluster binding (Figure 1A), but outside of that showed wide variations in conservation between their respective overall protein sequences. FDX1/2 were most similar among the group, at ∼80% similarity of their 94 (FDX1) or 93 (FDX2) amino acids (Figure 1A). While some Fd isoforms evolved additional C terminal domains, referred to as C-type Fds, FDX1/2 do not contain any C or N-terminal extensions. Where there are differences between FDX1/2, FDX2 is frequently adding additional charged amino acids compared to FDX1, 6 more charges in total, equally distributed between positively and negatively charged species (Figure 1A, changed charged amino acids are highlighted in red). The C-terminus contains the only position where FDX2 (S91) removes a charge present in FDX1 (E91), directly adjacent to a lysine (K92) in FDX2 where FDX1 has an alanine. Preceding this change is the only charge alteration between the two proteins, changing a histidine in FDX1 to an aspartate in FDX2 (H88D). These two changes combined effectively maintain the number of charges but invert the order of appearance, contributing to the difference in surface charge distribution between the two Fds(59, 86). Amino acids that contain an additional charge are among the most diverse positions between the different Chlamydomonas Fds, and amongst the group of FDX1/2 homologs in algae (Figure 1A, S1B). Most algae also encode for two Fds similar to FDX1/2 in Chlamydomonas, and while we do see differences between charged and uncharged amino acids in those positions between isoforms, outside of close relatives of Chlamydomonas they do not consistently distinguish the two isoforms over the larger evolutionary range, changes result in an opposite charges being added or are only used in a specific set of algae (Figure S1B). FDX2 was found to be more closely related to root-type Fds of plants (59), 4 of the 9 positions affected by those charge changes in Chlamydomonas are also different between leaf (FD1 and FD2) and root (FD3) type Fds of Arabidopsis. But unlike in Chlamydomonas, here more charges are found in the Fds predominantly expressed in photosynthetic tissues, not in root-type Fds (Figure 1A, green text). Taken together, this indicates that these positions are variable between similar Fds and are evolving separately in different species, likely to better accommodate individual client interactions.

The first three coordinating cysteines are in close proximity to each other (CX_4_CX_2_C), while the last cysteine regularly appears further downstream in the polypeptide chain. In case of FDX1 and FDX2, the same number of amino acids (29) are found between cysteine three and four, a similar protein architecture as is observed in other algae and plant homologs (Figure 1A, S1). A small subset of the Chlamydomonas Fd isoforms vary from the conserved 3 cysteine motif: while FDX6 has an additional Cys in the 3rd position in between the first two cysteines, FDX8 and FDX11 contain an additional, 5^th^ amino acid between the first two Cys (CX_5_CX_2_C). CX_5_CX_2_C has been commonly found in mitochondrial Fds (adrenodoxins) and, given that plastid localization has not been experimentally validated, current annotations may be erroneous (87). FDX8 and FDX11, however, are widely conserved in other algae, and distinct from algae adrenodoxins (Figure S1). Both proteins are conserved in marine and freshwater chlorophytes, showing distinct amino acid differences especially between and around the conserved cysteines involved in coordinating the [2Fe:2S] cluster. In algae adrenodoxins the charged amino acids here carry mostly negative charges, while FDX8/11 present both positive and negatively charged amino acids in those areas. FDX8 carries an arginine directly adjacent to first cysteine of the CX_5_CX_2_C group, a position that is consistently positively charged in chloroplast targeted Fds, while algae adrenodoxines have a conserved glutamate in the spot (Figure 1A, Figure S1A). FDX11 has a high similarity with NDF4 from *Arabidopsis thaliana*, which in this plant has been found to comigrate with the NADPH dehydrogenase (NDH) complex involved in cyclic electron flow (88). Together with the predicted chloroplast localization, this indicates that there may be indeed conserved CX_5_CX_2_C chloroplast Fds.

Taken together, we identified orthologs of most Chlamydomonas Fds in other eukaryotic algae (28) (Figure 1B, S1A). FDX2 shares high sequence similarity with FDX1 and does not harbor additional N or C terminal domains. We did notice, however, that FDX2 contains more charged amino acids as compared to FDX1, consistent with its previously determined surface charge differences (59).

### FDX1 and FDX2 stoichiometry changes when altering nitrogen sources

Chlamydomonas can assimilate nitrate and ammonium, but only nitrate assimilation requires a strictly Fd-dependent step. While FDX1 represents the major Fd isoform in Chlamydomonas, FDX2 is predominantly expressed when cells are grown on nitrate (NO_3_^−^) as the sole nitrogen source (59). Since it is unclear how much FDX2 contributes to the total Fd isoform protein pool, we set out to determine the relative stoichiometry between FDX1 and FDX2 at the protein level, when cells are grown on either nitrogen source (NH_4_^+^ and NO_3_^−^). Our approach to determine stoichiometry changes between FDX1 and FDX2 was to tag both proteins with the same immunoreactive epitope tag using a CRISPR based gene knock-in approach (89). To maintain native gene regulation, we modified the endogenous *FDX1* and *FDX2* gene loci using CRISPR/Cpf1 gene editing, providing single-stranded DNA (ssODN) templates. Our ssODNs contained homologous arms corresponding to the 3’ end of the FDX1 and FDX2 genes, respectively, and a short, codon-optimized sequence encoding a three glycine linker followed by an HA-tag (YPYDVPDYA) (Figure 2A, (90), intended to be inserted just before the stop codon at the respective gene locus (Figure 2A and Table 1). FDX1 and FDX2-specific antibodies had been raised previously using the most diverse, C-terminal regions of FDX1 and FDX2, respectively, as peptides for antibody production (59). We analyzed these Fd isoform specific antibodies together with a commercially available antibody raised against the HA tag to validate expression of the Fd-isoform-HA tag fusion proteins (Figure 2B, Figure S2B/C). Addition of the HA epitope to the C-terminus of FDX1, but not FDX2, resulted in a reduction of the affinity of the antiserum raised against Chlamydomonas FDX1 in detecting the FDX1-HA fusion protein (Figure S2C), which prompted us to add a polyclonal serum raised against spinach FDX1 to monitor FDX1 abundance in Chlamydomonas in parallel to the HA epitope (Figure 2B, S2C). As expected, we found that FDX1-HA and FDX2-HA fusion proteins showed no difference in band intensity but caused an increase in apparent molecular weight when separated on SDS-PAGE as compared to the native, untagged Fd isoform in wildtype cells (Figure 2B). No band was identified above detection limit at the unmodified size of either protein in any strains expressing HA-fusion proteins (Figure 2B). This data suggests that endogenous Fd isoform expression is not altered by the introduction of the linker and HA tag and that the respective Fd-HA fusion protein was the only protein produced from the edited, native locus. The strains expressing FDX1-HA or FDX2-HA fusion proteins did not have altered growth rates on photoheterotrophic or phototrophic growth media, either with NH_4_^+^ or NO_3_^−^ as nitrogen sources (Figure S2A). The antiserum against the HA epitope only recognized the modified FDX1-HA or FDX2-HA fusion proteins, respectively, and did not result in any signal in non-engineered wildtype samples (Figure 2B/C, Figure S2B).

**Figure 2.**
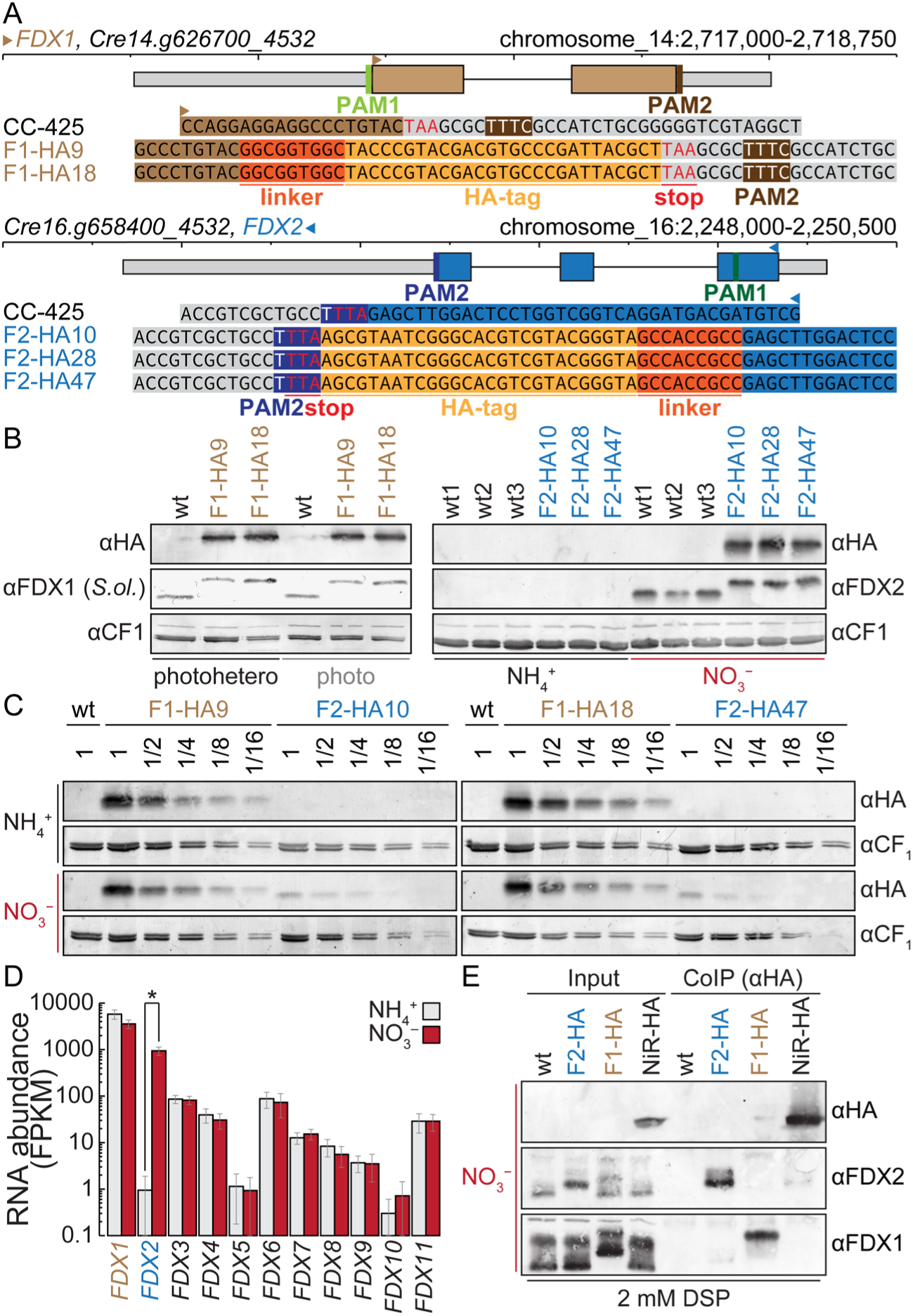
FDX2 is sub-stoichiometric to FDX1 on NO_3_ ^-^ but the preferred electron donor to NiR. (A) Illustration of the *FDX1* and *FDX2* gene models showing utilized PAM sites and introduced modifications to the different FDX1 (F1-HA1/2) and FDX2 (F2-HA1/2/3) lines carrying HA epitopes at their C-terminus. (B) Proteins were isolated from: (left) wildtype (wt) Chlamydomonas control and both FDX1 HA lines grown in phototrophic or photo-heterotrophic conditions with NH_4_^+^ as its sole N source; (right) wild-type controls and all three FDX2 HA lines grown photoheterotrophically on NH_4_^+^ or NO_3_^-^ as the sole nitrogen source. The proteins were separated with SDS-PAGE, transferred to nitrocellulose membranes and decorated with antibodies specific to FDX1/2 and the HA epitope. Antibodies specific to CF1 served as a loading control. (C) Protein isolated from FDX1 and FDX2-HA lines grown photo-heterotrophically on NH_4_^+^ or NO_3_^-^ were diluted and separated by SDS-PAGE, transferred to membranes and decorated with an antibody raised against the HA tag. CF1 serves as a loading control. (D) RNA abundance from wildtype Chlamydomonas, grown on NH_4_^+^ (grey) or NO_3_^-^ (red) as the sole nitrogen source was determined using RNAseq, error bars indicate standard deviation between three replicates. (E) Input material and co-immunoprecipitate (CoIP) from wildtype control, FDX2-HA, FDX1-HA and NiR-HA strain, separated by SDS-PAGE, transferred to membranes and decorated with an antibody raised against the HA tag, FDX1 and FDX2-specific antisera.

FDX1 abundance was unaffected by the presence or absence of acetate as an additional carbon source, both in wildtype and FDX1-HA-expressing strains (Figure 2B). Native regulation of the *FDX2* locus was maintained in strains expressing the FDX2-HA fusion protein, as FDX2 was only detected when NO_3_^−^ was provided as sole nitrogen source, either using the HA-specific or the FDX2 specific antibodies (Figure 2B). In nitrate grown cells, FDX2 accounted for ∼10% of the FDX1 abundance which is consistent with the magnitude of change observed at the transcript level (Figure 2C). Transcript profiling of cells propagated in the presence of nitrate or ammonium confirmed previous reports that FDX2 was indeed highly induced on nitrate and further showed that FDX2 is the only nitrate inducible Fd isoform in Chlamydomonas (Figure 2D).

### Nitrite reductase and FDX2, but not FDX1, interact *in vivo*

FDX2 is exclusively expressed on nitrate (Figure 2B) and had been shown *in vitro* to be more effective in catalyzing the NO_3_^−^ to NH_4_^+^ reduction, as compared to FDX1 (59). To identify which Fd interacts with NiR *in vivo*, we additionally introduced an HA-tag to the native 3’ end of the NiR gene (*NII1*, Cre09.g410750_4532), similarly to the approach used to generate strains expressing FDX-HA fusion proteins (Table 1). We identified two independent strains expressing NiR-HA fusion proteins (Figure S2B, Table 1) at the expected size that we utilized alongside wildtype, FDX1-HA and FDX2-HA expressing strains to purify complexes from cell lysates using HA antibodies via Co-Immunoprecipitation (CoIP, Figure 2E). We split the purified complexes for protein identification using both targeted analyses by immunodetection and untargeted detection via mass-spectrometry. This was necessary to recover FDX1, which we failed to detect in any MS-based approach but consistently recovered in immunodetections, while FDX2 was recovered only once in MS (Figure 2E, Figure S2D, Data S2). We were unsuccessful in establishing a reciprocal Fd-client interaction without the use of a chemical crosslinker with either approach but determined reciprocal FDX2-NiR interaction when using 2 mM Dithiobis(succinimidyl propionate) (DSP) to stabilize complexes (Data S2, Figure 2E). While NiR was consistently recovered in precipitates from FDX2-HA strains, even without crosslinker, we did not identify NiR in FDX1-HA samples (Data S2). FDX2, but again not FDX1, was enriched in precipitates from crosslinked NiR-HA tagged lines in immunoblots (Figure 2E). In all analyses, we never identified FDX1, FDX2 or NiR in precipitates from wild-type controls, indicating that our immunoprecipitations using the HA-tag resulted in specific enrichment (Data S2).

While both IP-MS and IP-immunodetections have limitations in sensitivity, and the stoichiometry between FDX1:FDX2 of ∼10:1 in nitrate grown cells is in favor of FDX1, these results are in line with earlier studies proposing that FDX2 is indeed the preferred electron donor to NiR *in vivo*.

### Chlamydomonas strains with a complete loss of FDX2 can assimilate NO_3_^−^

Since the kinetics and higher *in vivo* binding affinities suggested a role for FDX2 in nitrite reduction, we hypothesized that FDX2 is required for algal growth on medium containing NO_3_^−^ as the sole nitrogen source. To test this, we generated *fdx2* mutants using CRISPR/Cpf1 gene editing, this time, introducing *inframe* stop codons within the first exon of *FDX2* (Table 1), resulting in early termination 8 amino acids into the mature FDX2 protein (Figure 3A). In this case we used diagnostic PCR to identify mutants that contain perfect stop codon insertions, and genotypes were subsequently confirmed with Sanger or whole genome sequencing. We found three independent strains, depicted *fdx2-1*, *fdx2-2* and *fdx2-3,* that do not express FDX2 on NO_3_^−^ (Figure 3 B/C). FDX2 protein was not detectable in all three *fdx2* mutants, with FDX2 expression seemingly unaffected by carbon availability (Figure 3C). Interestingly, *FDX2* transcripts did also not accumulate in *fdx2* mutants on NO_3_^−^ (Figure 3B), suggesting that the addition of early *in-frame* stop codons resulted in nonsense-mediated mRNA decay. To test the contribution of FDX2 to N assimilation we grew *fdx2* mutants alongside wild-type controls with either NH_4_^+^ or NO_3_^−^ as the sole N source, both in phototrophic and photoheterotrophic conditions (Figure 3D). The pressure to utilize FDX2 for electron delivery to NiR should be higher in phototrophically grown cells, as they also depend on photosynthetically-driven carbon assimilation via Fds for growth. Surprisingly, *fdx2* mutants grew well, with similar growth rates compared to wild-type strains and on either N or C source (Figure 3D). We then probed expression of genes encoding proteins involved in N assimilation using RNA-seq and qRT-PCR and found that transcript abundance was largely unchanged between wildtype and *fdx2* mutants (Figure 3E, Figure S3A). Most transcripts involved in NO_3_^−^ assimilation accumulated to similar levels in wild-type strains and *fdx2* mutants, as were those for high-affinity NH_4_^+^ transporters and transporters/assimilatory proteins involved in the assimilation of other, lower priority N sources (Figure 3E). While the vast majority of genes coding for central components involved in nitrogen assimilation were expressed normally, we noticed reduced accumulation of two transcripts in *fdx2* mutants: *NRT2C* and *LAO2*. *NRT2C* encodes for a nitrite importer into the cytosol and *LAO2* codes for an accessory protein to periplasmic L-amino acid oxidase, both proteins are involved in N assimilation but are not be required for growth on nitrate. Both genes were induced on NO_3_^−^ in both wildtype and *fdx2*, but the amplitude of induction was ∼2 fold lower in *fdx2* mutants (Figure 3E, Figure S3A).

**Figure 3.**
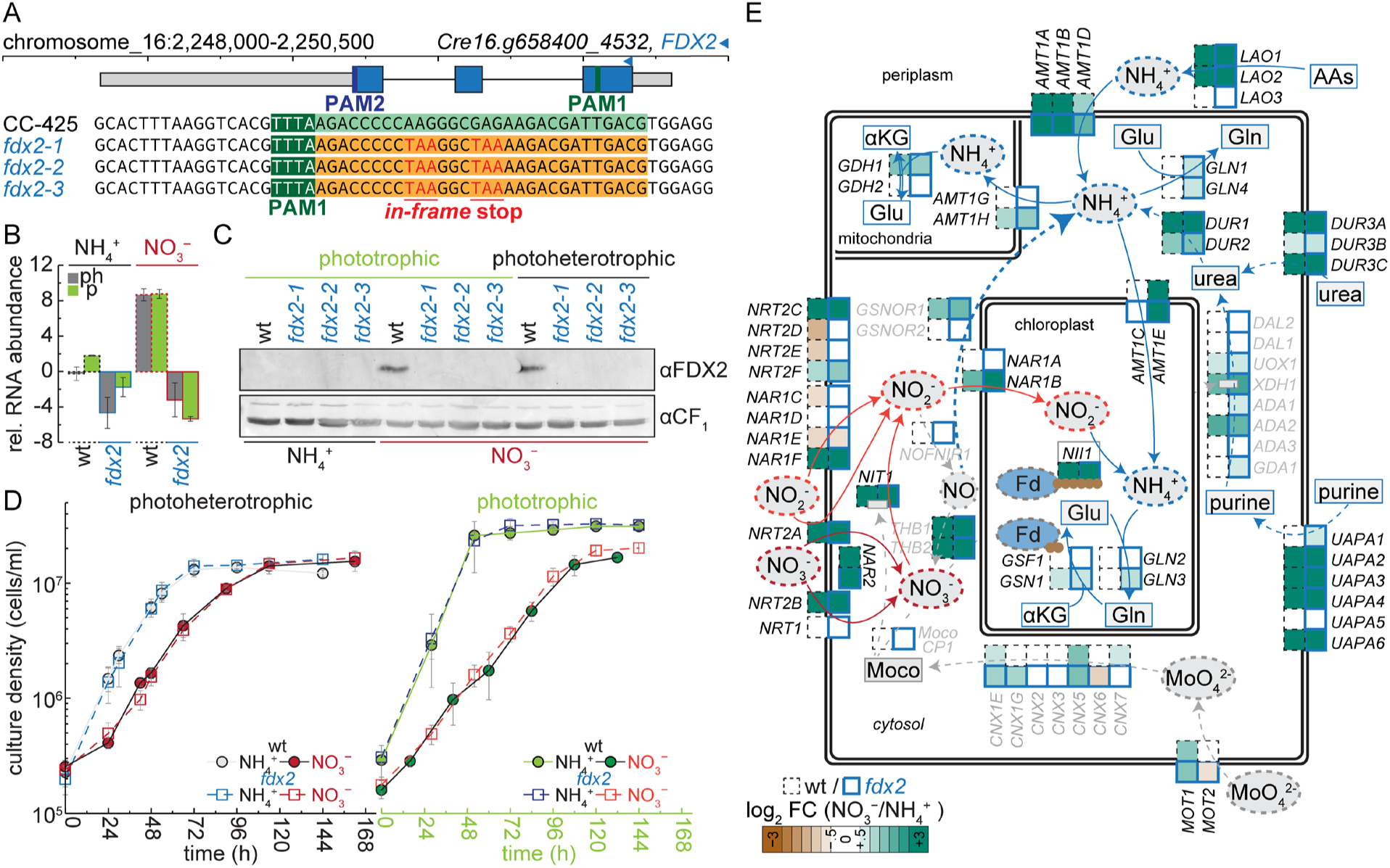
*fdx2* knockouts are phenotypically wildtype when grown on nitrate. (A) FDX2 locus, utilized PAM sequences for Cpf1 recruitment and ssODN-mediated gene editing, resulting in the insertion of two *in-frame* stop codons into exon 1. (B) qRT-PCR assessment of *FDX2* transcript abundance in RNA extracted from *fdx2-1*, *fdx2-2* and *fdx2-3* grown alongside a wildtype controls photoheterotrophically (grey fill) or phototrophically (green fill) on NH_4_^+^ or NO_3_^-^ as the sole nitrogen source. Shown are the averages and standard deviation from three independent experiments. (C) Whole cell protein from cultures was separated by 15% SDS-PAGE, transferred to nitrocellulose membranes and immunodecorated with antisera against FDX2. CF1 was used as loading control. (D) Growth of wildtype (circles) and *fdx2* (squares) in various conditions, shown are averages and standard deviation from three independent wildtype strains or *fdx2* mutants. (E) Transcript abundance changes between NH_4_^+^ and NO_3_^-^ (log fold change) in wildtype (black, dashed outline) and *fdx2* (blue, solid outline) for genes involved in nitrogen assimilation. Blue fill indicates increasing transcript abundance on NO_3_^-^, brown fill decreasing abundance, shown is the average of 3 wildtype or *fdx2* strains.

This data indicates that, consistent with the lack of a growth phenotype, nitrate assimilation was not negatively impacted by a loss in FDX2 function. Taken together we conclude that FDX2 is not essential for growth on NO_3_^−^, and a different Fd, most likely FDX1, can supply electrons efficiently to NiR and completely compensate for the loss of FDX2 function.

### *fdx1* mutants grow better with NO_3_^−^ as N source

To dissect the potential, partial functional redundancy between FDX1 and FDX2, we aimed to generate *fdx1* mutants. Our initial attempts to generate CRISPR-based *fdx1* knockouts by introducing *in-frame* stop codons early within the *FDX1* coding sequence, similar to the strategy used for *FDX2*, were unsuccessful (data not shown). We reasoned that FDX1 may be essential and aimed for an inducible knock-out strategy instead. In Chlamydomonas, thiamine biosynthesis is controlled by riboswitches, which are regulatory, non-coding RNAs. Two transcripts coding for enzymes required for early steps in thiamine biosynthesis contain characterized riboswitches: *THI4* and *THIC* (91, 92). The *THIC* riboswitch affects splicing of an intron within the gene body, resulting in an early termination event. The *THI4* riboswitch is in the 5’UTR, also involving introns facilitating an alternative splicing event, but in this case resulting in the translation of a µORF instead of the main ORF when either thiamine or thiamine pyrophosphate (TPP) are present in the growth media (91). This mechanism allows Chlamydomonas and many other eukaryotes to avoid costly synthesis of thiamine when they find it in their environment. To be able to conditionally repress *FDX1* expression, we used single stranded DNA templates (ssODNs) containing homology arms corresponding to the *FDX1* 5’ region adjacent to the start codon (ATG), as well as the 1114 bp Chlamydomonas *THI4* 5’UTR including the riboswitch (Figure 4A). If this system works as intended, FDX1 protein should be conditionally depleted by growing cells in the presence of thiamine, expressing the µORF instead, resulting in inducible *fdx1* null alleles. We screened 138 transformants for their ability to grow on growth medium that is supplemented with 10 µM thiamine and identified one strain, *fdx1-rs*, in which we observed a thiamine-dependent growth defect. Whole genome sequencing of *fdx1-rs* showed that the full *THI4* riboswitch sequence was integrated upstream of the start codon, while a small, 7 nt sequence of the native FDX1 5’UTR just upstream of the PAM site used for integration was removed (Figure 4A). *FDX1* mRNA abundance in *fdx1-rs* was reduced already in the absence of thiamine, ∼ 10x compared to levels found in wild-type (Figure 4B). This may be attributed to the changes made to the native *FDX1* 5’UTR, which supports the high expression of this Fd. While wild-type *FDX1* mRNA levels remained unchanged regardless of thiamine in the growth medium, the addition of thiamine to *fdx1-rs* mutant resulted in a further decrease in *FDX1* transcript levels (Figure 4B, left panel). Beyond *fdx1-rs*, we identified several lines exhibiting thiamine-independent growth defects (termed *fdx1-1/2/3,* Figure 4C, Figure S4A/B/C). We used whole genome sequencing to identify the modifications made to the *FDX1* locus in these strains and found the coding region to be unaffected in all of them, but the 5’UTR to be modified, either by incomplete integration of the *THI4* riboswitch or other sequences. *fdx1-1* has a 33nt deletion of the native *FDX1* 5’ UTR, which was replaced by a 614nt insertion that contains potential ORFs. *FDX1* transcripts in *fdx1-1* were similarly reduced than in *fdx1-rs*, ∼10x compared to wildtype, which was unaffected by thiamine presence (Figure 4B, S4C). Native *THI4* mRNA levels remained consistent across wildtype and all *fdx1* mutants, *THI4* mRNA abundance was also reduced with thiamine addition (Figure 4B, middle panel). The nitrogen source (NH_4_^+^ or NO_3_^−^) did not affect expression of *FDX1* or *THI4* expression, *NIT1*, encoding for nitrate reductase was used a marker to assess nitrogen source use, was also induced similarly in all strains on NO_3_^−^, independent of the presence of thiamine (Figure 4B, right panel). At the protein level, *fdx1-rs* showed a severe reduction of FDX1 to between 2 and 5% of wild-type levels, which was further reduced to non-detectable levels ∼ 72 hours after the addition thiamine (Figure 4C). *fdx1-1/2/3* had even further reduced FDX1 protein amounts, below 1% of wild-type abundance (Figure 4C). FDX1 abundance was unchanged when cells were grown on different nitrogen sources, as was the abundance of FDX2 in *fdx1-rs* strains that grew on nitrate (Figure 4C, right panel).

**Figure 4.**
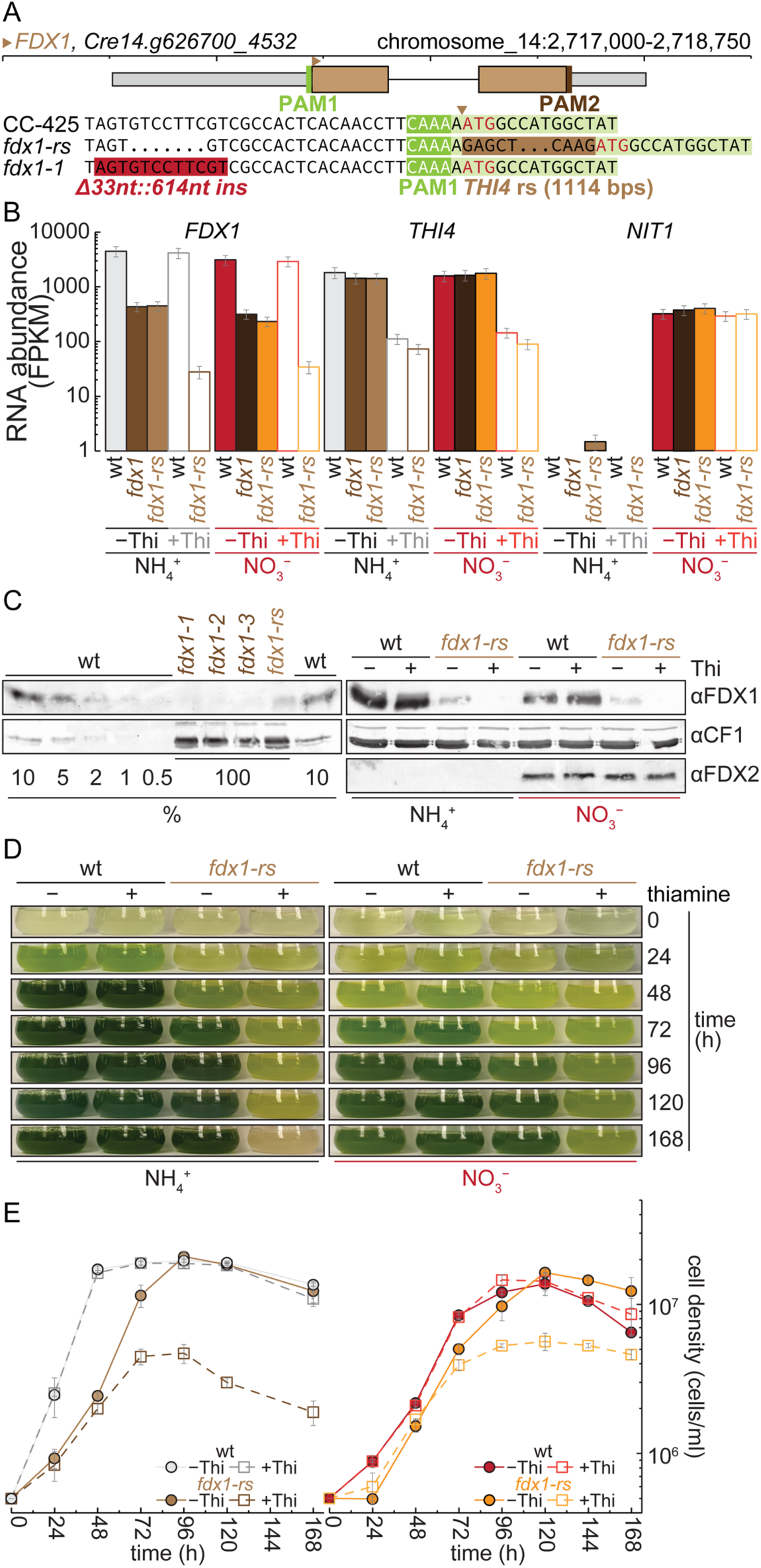
*fdx1* (inducible) knock out strains are non-viable. (A) Physical map of the *FDX1* gene and PAM sequence for Cpf1 recruitment. CRISPR/Cpf1-and ssODN-mediated gene editing resulted in a 614 bp insertion in *fdx1-1* or the insertion of the *THI4* riboswitch in *fdx1*-rs. (B) Transcript abundance of *FDX1*, *THI4* and *NIT1* in RNA extracts from wildtype, *fdx1-1* and *fdx1-rs* grown photo-heterotrophically on NH_4_^+^ or NO_3_^-^, with or without thiamine, as indicated. Transcript abundance was analyzed by RNAseq, normalized to gene length and sequencing depth (fpkm), shown are average and standard deviation of three independent cultures. (C) Whole cell protein extracted from wildtype and various *fdx1* knock-down strains was separated by 15% SDS-PAGE, transferred to nitrocellulose membranes immune-decorated with anti-sera against FDX1 or FDX2, respectively. CF_1_ was used as loading control. (D/E) Growth of wildtype and *fdx1-rs* on NH_4_^+^ or NO_3_^-^, with or without thiamine, as indicated, shown are average and standard deviation of 3 independent cultures.

While we already noticed growth defects during the screening process, we systematically analyzed and quantified growth of the individual *fdx1* mutants further. Interestingly, FDX1 appears to have an essential role in photoheterotrophic conditions, where cells are less reliant on Fd-dependent photosynthetic electron distribution, especially towards the major sink CO_2_ fixation. *fdx1* knockdowns that contained less than 1% residual FDX1 (*fdx1-1*) showed reduced growth rates on both nitrogen sources but were eventually able to reach cell densities similar to wildtype (Figure S4B). We did not observe growth differences in wildtype or *fdx1-1/2/3* with respect to thiamine supply. *fdx1-rs* cells that contained ∼ 2-5% of residual FDX1 (no thiamine addition), showed a similar reduction in growth rate as *fdx1-1* on NH_4_^+^ compared to wildtype, but were less affected on NO_3_^−^ where FDX2 expression was induced (Figure 4DE, Figure S4E). Thiamine-induced *fdx1-rs* cultures cede to grow after 2-5 days after thiamine addition, depending on carbon and nitrogen source (Figure 4C, Figure S4E). 5 days after thiamine supplementation, *fdx1-rs* cells grown on NH_4_^+^ started dying as cell numbers declined, which was not observed in NO_3_^−^ grown cultures (Figure 4C). *fdx1-rs* cells also reached higher cell densities at peak (Figure 4E), and similar growth rates to wildtype early after thiamine addition on NO_3_^−^ (Figure 4E, Figure S4C). The nutritional rescue of *fdx1-rs* null mutants (+ thiamine) on nitrate suggested that FDX2 may partially compensate for the absence of FDX1 function (Figure 4D). As expected, induced *fdx1-rs* strains were also non-viable in phototrophic growth media when thiamine was added, while *fdx1* knockdowns (*fdx1-1*) and uninduced *fdx1-rs* still managed to reach stationary cell densities, albeit at reduced growth rates compared to wildtype (Figure S4E). Chlorophyll content was comparable between *fdx1*, *fdx2* mutants and wildtype controls, also implying that a lack of coloration did not result from a lack in pigmentation, but rather was the result of stunted growth (Figure S5B).

Taken together, between 1-5% residual FDX1 is sufficient to almost completely rescue the lethal phenotype of *fdx1* null mutants. While the degree of an observable growth phenotype scaled with the level of FDX1 reduction, *fdx1* culture growth improved on nitrate, where FDX2 is expressed.

### FDX1 and FDX2 together are essential for growth on NO_3_^−^

The observation that neither the *fdx1* nor the *fdx2* single mutants have an exacerbated phenotype on NO_3_^−^ led us to the hypothesis that the presence of either Fd is adequate to accommodate NO_3_^−^ assimilation. To test this, we attempted to generate *fdx2* knockout strains in the *fdx1-rs* background, using the same PAM site and ssODN that were utilized to generate the *fdx2* knockouts in the first place (Figure 5A, Table 1). For selection after transformation we used paromomycin resistance, instead of restoring arginine auxotrophy, which was utilized to generate *fdx1-rs, fdx1-1* and wildtype controls of the recipient strain. We identified two strains that again successfully added two *in-frame* stop codons into the first exon of *FDX2*, named *fdx1-rs fdx2-1* and *fdx1-rs fdx2-2*, with *fdx1-rs fdx2-2* harboring an additional 3bp mutation downstream of the gene editing site (Figure 5A). While *fdx1-rs* still accumulated FDX2 on NO_3_^−^, *fdx1-rs fdx2* mutants, similar to *fdx2* single knockouts, had no detectable FDX2 protein left (Figure 5B). Without thiamine addition, *fdx1-rs* strains carrying the stop codons in the first exon of *FDX2* behave like *fdx1-rs* on NH_4_^+^, which was not surprising since FDX2 is not expressed when NH_4_^+^ is present in the growth media (Figure 5C). On NO_3_^−^, in the presence of thiamine, *fdx1-rs fdx2* fail to grow almost immediately after thiamine addition (Figure 5C).

**Figure 5.**
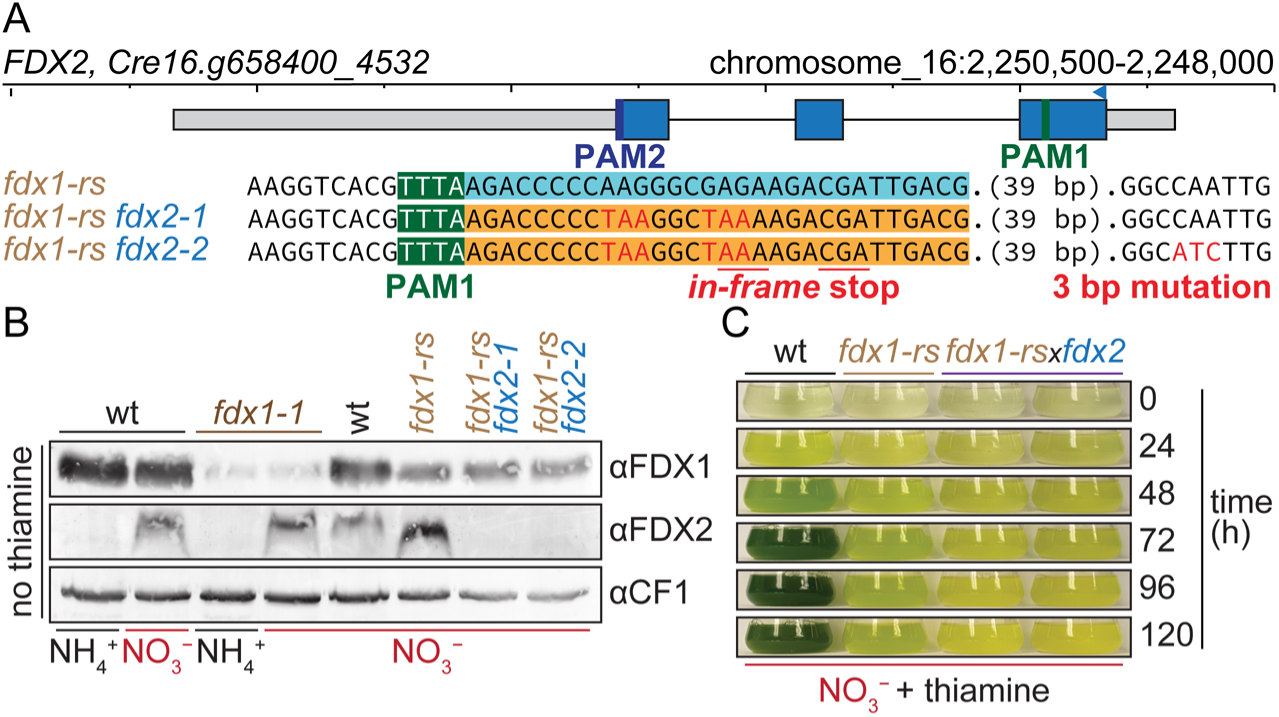
FDX1 or FDX2 are required for growth on NO_3_^-^. (A) *FDX2* locus in *fdx1-rs, fdx1-rs fdx2-1/2*, utilized PAM sequences for Cpf1 recruitment and ssODN-mediated gene editing, resulting in the insertion of two *in-frame* stop codons into exon 1 of FDX2. (B) Whole cell protein extracts from wildtype, *fdx1-1*, *fdx1-rs, fdx1-rs fdx2-1/2* cultures, grown photoheterotrophically on NH_4_^+^ or NO_3_^-^ without thiamine, were separated on 15% SDS-PAGE, transferred to nitrocellulose and immunodecorated with antibodies specific for FDX1 and FDX2, CF_1_ served as loading control. (C) Growth on NO_3_^-^ in the presence of thiamine was monitored by taking pictures of flasks. Shown are representative pictures of three independent experiments.

From this we conclude that FDX1 is indeed the Fd substituting for FDX2 in *fdx2* mutants, facilitating growth on NO_3_^−^. We also conclude that FDX1 or FDX2 alone are sufficient to support growth on media utilizing NO_3_^−^ as the sole N-source, but in their absence no other Fd can substitute for this function.

### *fdx2* is affected only in low abundance, auxiliary processes for C, N and S assimilation

Both, *fdx1* knockdowns and knockouts, exhibited (severe) growth defects on both N sources, while *fdx2* mutants were largely unaffected. To determine which aspect of Fd-dependent metabolism was mis-regulated in the respective mutants, we conducted three independent RNA-seq experiments comparing control cultures to *fdx1-1*, *fdx1-rs* (+/- thiamine) or *fdx2* strains (Figure 6A, Data S3). The wildtype and mutant strains were grown in either NH_4_^+^ or NO_3_^−^ media in photoheterotrophic conditions. All cultures were prepared in biological triplicates and collected at mid-log cell density for analysis, for thiamine-induced *fdx1-rs* ∼ one doubling prior to growth arrest (Figure 6A and Figure S6). In a principal component analysis, the individual replicates grouped well based on genotype/condition, with expression differences between NH_4_^+^ and NO_3_^−^ contributing strongly to the separation of the samples in all comparisons (Figure 6A, Figure S6E). This was most pronounced for *fdx2* mutants, but less so in *fdx1* strains, correlating with decreasing FDX1 abundance (PC1, Figure 6A). The effect of the mutations was found in the 2^nd^ principal component, where it was more impactful for *fdx1-1* (∼25% of variance) than for *fdx2* (∼16% of variance, Figure 6A). For *fdx1-rs* sample separation was more equally driven by mutation and N source, and we noticed a clear difference between thiamine-induced and uninduced *fdx1-rs* mutant stains, while wildtype samples did not show separation depending on thiamine supplementation, both on NH_4_^+^ or NO_3_^−^ (Figure 6A, middle panel).

**Figure 6.**
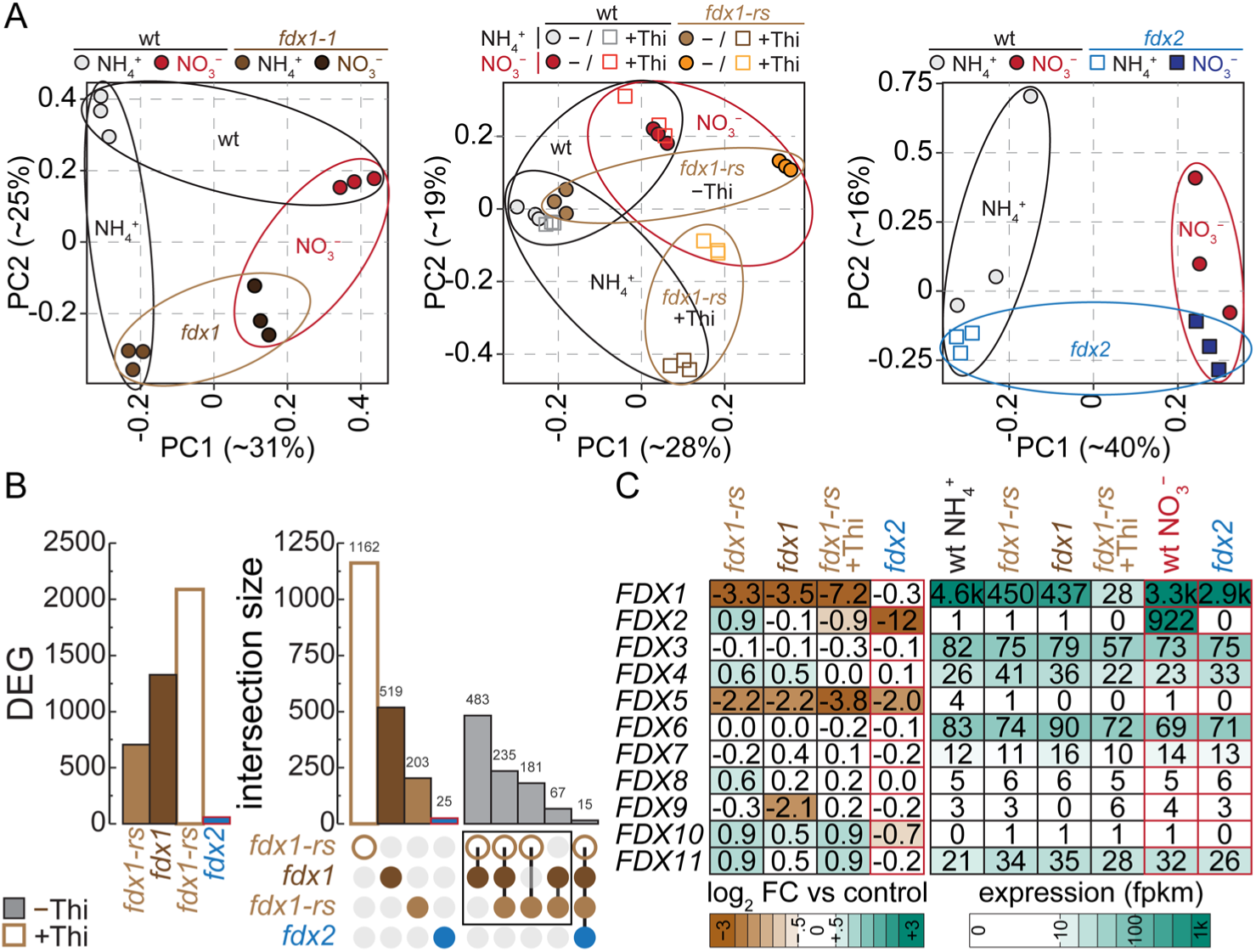
Transcriptome changes to FDX1/2 depletion. (A) Principal component analysis of the 3 RNAseq experiments. Shown are the individual samples of wildtype controls, *fdx1-1*, *fdx1-rs* and *fdx2* mutants grown photoheterotrophically either on NH_4_^+^ or NO_3_^-^, with or without thiamine. (B) Number of differentially expressed genes (DEG, left panel), de-fined as at least a 2-fold change in abundance (5% FDR), between the respective control and indicated mutant. Red outlines indicate NO_3_^-^ growth conditions, filled bars indicate no thia-mine addition. UpSet (right panel) highlighting unique changes to individual treatments as well as subsets of overlap. Red outlines indicate NO_3_^-^ growth conditions, filled bars/ circles indicate no thiamine addition. (C) Heatmap showing log_2_ fold changes of transcript abundances of chloroplast Fds between respective control and indicated mutant (left), brown indicates lower abundance in the mutants, blue indicates increased abundance, and average transcript abundance (right, fpkm), darker indicates higher abundance. Red outlines indicate NO_3_^-^ growth conditions.

We used a log_2_ fold change cutoff of ±1 and an adjusted p-value of 0.05 based on the variance between replicate cultures to identify differentially expressed genes (DEG) between different strains and conditions for all experiments (Data S4). We found very few changes between wildtype and *fdx2* mutants grown both with NH_4_^+^ (160 genes) or NO_3_^−^ (59 genes) as the only N source (Figure 6B, Data S4). Transcripts of other chloroplast-target Fds were not induced in *fdx2*, indicating that the absence of FDX2 can largely be compensated for by the existing Fd pool (Figure 6C). The majority of differentially expressed genes were expressed at low levels in wildtype and most of them showed a mild reduction in transcript abundance in *fdx2* (Data S4). *FDX2* was the most reduced gene on both N sources, albeit being barely expressed on NH_4_^+^ (< 1 fpkm, Figure S3, Data S3/4). FDX5 was the only other Fd that showed a significant transcriptional response; it was found to be reduced in the *fdx2* mutants when grown on NO_3_^−^ (Figure 6C). While previous studies found that the conditionally expressed *FDX5* is highly induced by anaerobiosis and copper deficiency (93, 94), it was expressed at very low levels in the aerobic, copper-replete samples analyzed here (around 1 fpkm in wild-type). Interestingly, 21 of the 30 genes whose transcripts were found to be most significantly reduced in *fdx2* mutants on NH_4_^+^ were increasingly accumulating on NO_3_^−^ both in wt and *fdx2* mutants, including low expressed genes in N assimilation (Data S4). The induction of these genes between NH_4_^+^ and NO_3_^−^ was more pronounced in *fdx2* mutants compared to wt controls, resulting in comparable levels of transcripts on NO_3_^−^ despite a significant reduction on NH_4_^+^ (Figure 3E, Data S4). We did find genes involved in N (*AMT1/2*, *NRT2C*, *AMX1*, *XUV1/5*), S (*ARS1/2*) and C (*PCYA*, *CAH1/4/5*, *HLA3*, *LCIA*) assimilation being less expressed in *fdx2* on NH_4_^+^. While genes involved in nitrate assimilation were not differentially expressed in *fdx2*, genes coding for high affinity ammonium transporters (*AMT1/2*), which are induced when ammonium becomes scarce (95–97), amine oxidase and transporters for other, lower priority N sources were reduced (*XUV1/5*) (Data S4). We did not notice consistent discrepancies in the expression of the more central components involved in these pathways, specifically respiration (Figure S7), chlorophyll biosynthesis and photosynthesis (Figure S5), S and NH_4_^+^/NO_3_^−^ assimilation (Figure 3E, Figure S3). For example, central genes to NO_3_^−^ assimilation showed basal expression on NH_4_^+^ in controls, still very low compared to their expression on NO_3_^−^, including nitrate (*NIA1*, ∼5 fpkm) and nitrite reductase (*NII1*, ∼1 fpkm), transporters in the plasma membrane (*NAR1C/D/E*) and chloroplast envelope (*NAR1B*), encoding a set of proteins that would allow for efficient import and assimilation of nitrate/nitrite in the chloroplast (Figure 3E, S3).

### *fdx1* knockdown already severely affects expression

In contrast to *fdx2* mutants, changes to the transcriptomes of *fdx1* knockdown or knockout strains were more impactful, with the extent scaling with the degree of *FDX1* removal (Figure 6B). In ammonium-grown *fdx1-rs* cells, where FDX1 levels were reduced to between 2% and 5% of wild-type levels, accumulation of 700 transcripts was significantly altered (Figure 6B, 4C). In the case of *fdx1-1* (0.5 - 1 % FDX1 remaining) we saw changes to ∼1300 transcripts, while in thiamine-induced *fdx1-rs* (< 0.1 % FDX1) ∼2100 genes were differentially expressed (Figure 6B). There was a significant overlap between the individual *fdx1* mutants, most of the genes changing in the least FDX1-reduced strain (thiamine-uninduced *fdx1-rs*,∼ 500 of 700 genes) were similarly affected in the other knockdowns, while thiamine-induced *fdx1-rs* had the most unique changes, but still about half of the genes changing were found similarly changing in the knockdowns (Figure 6B). Similar to what we observed in *fdx2* mutants, *fdx1* mutants did not show any significant change in expression of other chloroplast-targeted Fds (Figure 6C). Since all *fdx1* mutants exhibited a growth defect (Figure 4DE, Figure S4), this indicates that while the abundance of individual Fds can be adjusted transcriptionally, there is no specific transcriptional regulatory program controlling Fd expression when FDX1 or FDX2 are absent. Individual Fds seem to be transcriptionally regulated within the metabolic modules in which they participate (for example FDX2 on NO_3_^−^, Figure 2D).

While *fdx2* mutants did not show differential expression of genes involved in N assimilation (Figure 3C, Figure S3), nitrogen assimilation was affected in the different *fdx1* knockdowns (Figure S8). N assimilation was more prominently changed in cultures grown on NO_3_^−^, and primarily affected transporters for lower priority N sources. These transporters were reduced in expression, including the conditionally induced NO_3_^−^ importers (*NRT2C*, *NAR1B/F*), high affinity NH_4_^+^ transporters (*AMT1A/B*) and urea importers (*DUR3A/C*). L-amino acid oxidase 1 (*LAO1*) was induced in *fdx1-1* and thiamine-induced *fdx1-rs*, but not in thiamine-uninduced *fdx1-rs* where 2-5% FDX1 was left (Figure S8). Expression of the lesser expressed of two truncated hemoglobins (THB1) and nitrosoglutathione reductase (GSNOR1) were reduced consistently in *fdx1* mutants, both of which are involved in nitric oxid (NO) processing originating from nitrate reductase. Expression of GS/GOGAT, nitrate and nitrite reductase, many of the highest expressed transporters for NO_3_^−^/ NO_3_^−^/NH_4_^+^ (*NRT2A*, *NAR1A*, *NAR2*, *AMT1D*), and more importantly genes facilitating the assimilation of even lower priority N sources were not differentially accumulating (Figure S8).

Taken together, our transcriptome data suggests that *fdx1* mutants still assimilated NO_3_^−^ but changes to gene expression would accommodate a reduced flux in N assimilation, commensurate with slower growth rates.

### *fdx1* mutants are S deficient

One prominent aspect of Fd-dependent metabolism affected in *fdx1* mutants was S assimilation (Figure 7A/B). Chloroplast sulfite reductases require electrons from Fds to catalyze the 6-electron reduction of sulfite to sulfide. Unlike for N assimilation or photosynthesis, gene expression for S assimilation was mostly increased and involved the most highly expressed isoforms of the individual transporters or enzymes. On NH_4_^+^, sulfate importers *SLT1/2/3* and *SULTR2* were found to be significantly induced in *fdx1* knockdowns; the extent of the increase scaled with the amount of FDX1 reduction (Figure 7A/B). Chloroplast importer *SULP3* was increasingly expressed in all *fdx1* mutants except thiamine-uninduced *fdx1-rs*. Chloroplast enzymes involved in S assimilation into cysteine (*ATS1*, *SIR1/2*, *SAT1*, *OASTL4*), all of which were the predominantly expressed isoform in wildtype controls, were found to be increased as well (Figure 7A/B). Among the four genes coding for O-acetylserine (thiol)lyases in Chlamydomonas, *OASTL4* was also the only induced gene in S-deficiency (72). Some additional genes, found to be induced in response to S-deficiency, aryl sulfatases *ARS1/2*, cysteine dioxygenase *CDO1*, taurine/alpha-ketoglutarate dioxygenases *TAUD1/2*, rhodanese *RDP3*, low S extracellular cell wall proteins *ECP56/76/88*, were also induced in *fdx1* knockdowns, for most of them, the induction again scaled with the amount of FDX1 protein reduction. These changes in gene expression indicated a lack of S and the desire to increase its assimilation. We therefore measured the S content in controls, *fdx2* mutants, *fdx1-rs* knockdowns with or without thiamine supplementation using ICP MS/MS (Figure 7C). Despite the increase in gene expression, cellular S levels in *fdx1-rs* were significantly reduced on NH_4_^+^ in thiamine uninduced *fdx1-rs* (Figure 7C). The S content decreased further in thiamine induced *fdx1-rs*, where the expression of more genes involved in S assimilation was increased (Figure 7A/B/C).

**Figure 7.**
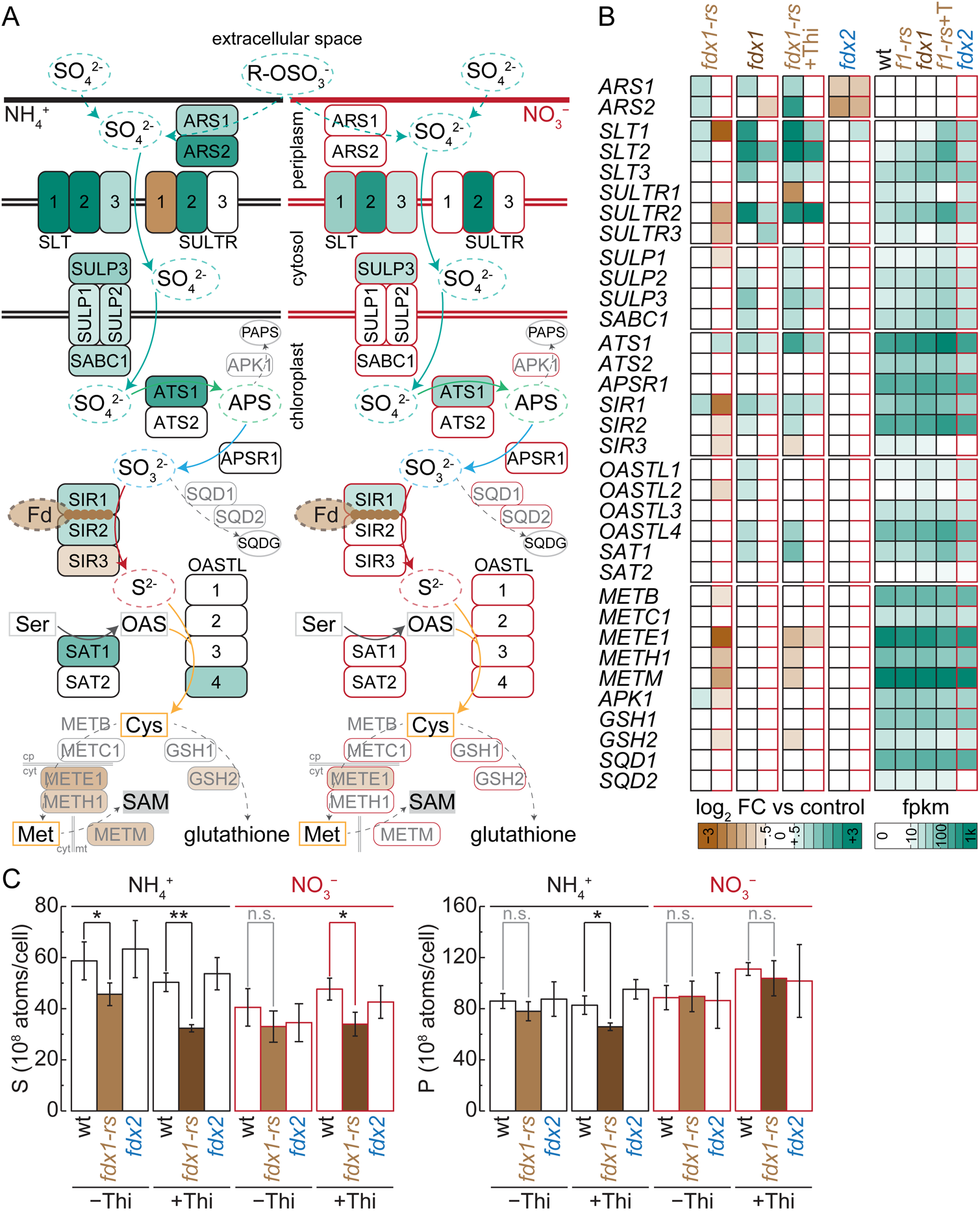
Sulfur assimilation is increasingly induced with less FDX1 presence. (A) Overview of proteins involved in sulfur assimilation in *C. reinhardtii*. Each fill color reflects the differences (log_2_ fold changes) of transcript abundance between wildtype and *fdx1-rs* (thiamine induced), brown indicates lower abundance in the mutants, blue increased transcript abundance, either on NH_4_^+^ (left, black outlines) or NO_3_^-^ (right, red outlines). (B) Heatmap showing the full summary of log_2_ fold changes of transcript abundances between respective wildtype controls and indicated mutant (left), brown indicates lower abundance in the mutants, blue indicates increased abundance, and average transcript abundance (right, fpkm), darker indicates higher abundance. Red outlines indicate NO_3_ ^-^ growth conditions. (C) Sulfur and phosphorus content of wildtype, *fdx1-rs* and fdx2 mutants, measured by ICP MS/ MS, shown are average and standard deviation of three independent cultures. Asterisks indicate significant changes (* p < 0.05, ** p < 0.01).

*fdx1*-rs mutants were more viable when grown on nitrate where FDX2 was expressed (Figure 4D), concomitant with a reduced induction of the S deficiency response (Figure 7A/B). We therefore wondered if nitrate grown *fdx1* mutants can maintain their S quota. On NO_3_^−^, most of the genes induced on NH_4_^+^ were no longer significantly increased or showed lower magnitudes of induction, in some cases even reduced expression compared to wildtype controls (Figure 7A/B). Thiamine-uninduced *fdx1-rs* fared the best, which coincided with wildtype levels of cellular S (Figure 7C), while in thiamine-induced *fdx1-rs* most genes in S-assimilation were still induced and S-content was still reduced compared to wildtype controls. *fdx2* mutants did not show any changes to S assimilation or cellular S content (Figure 7B/C).

While transcripts for proteins involved in S assimilation were induced, downstream of cysteine, genes encoding enzymes involved in methionine biosynthesis (*METE1*, *METM*) were significantly reduced in thiamine-induced *fdx1-rs* on NH_4_^+^, which was also found less pronounced on NO_3_^−^ (Figure 7A/B). Among the identified regulatory proteins involved in mediating the alga’s response to S deficiency, expression of kinase SNRK2B, a repressor of the S-deficiency response, was reduced in all *fdx1* knockdowns, significantly only in thiamine-induced *fdx1-rs* (Data S5). Expression of transcription factor Ars73a was also found to be induced in thiamine-induced *fdx1-rs*.

Taken together we conclude that FDX1 is required for S-assimilation, while cultures on NO_3_^−^, where FDX2 was expressed, were able to assimilate S better. Some residual amounts of FDX1 in combination with the expression of FDX2 were sufficient to sustain S assimilation, while further reduction of FDX1 still resulted in altered expression and lower cellular S.

### Fe assimilation and Fe-S cluster biogenesis are affected by fdx1 depletion

S is a key component of *de-novo* Fe-S cluster assembly in the chloroplast via the SUF system (98). The first step of this process in the chloroplast is the desulfuration of cysteine, catalyzed by a cysteine desulfurase (SufS/NFS2), forming a persulfide group on a catalytic cysteine. The bound S is handed over to the SUFE sulfur relay which donates the S to the Fe/S cluster scaffold protein SufB, a part of a tetrameric complex (SUFBC_2_D). In thiamine-induced *fdx1-rs* we found that both a homolog of NFS2 and SUFE were significantly induced, while only NFS2 was already increased in *fdx1-1* knockdowns. Downstream components of the SUF Fe-S cluster assembly machinery were not changed in their expression, including the scaffolding proteins themselves, which were expressed similarly in all different *fdx1* knockdowns (Figure S9).

Fds are abundant Fe proteins utilizing [2Fe-2S] clusters for electron transport, which prompted us to analyze the expression of genes involved in Fe assimilation. The Chlamydomonas genome supports both Fe(II) and Fe(III) assimilation, encoding for an Fe reductase (FRE1) and two ZIP family Fe(II) importers (IRT1/2) as well as a plasma membrane-bound Cu-dependent Fe oxidase (FOX1) that supplies Fe(III) to an Fe(III) permease (FTR1) (99, 100). These import systems are further supported by two highly abundant, excreted periplasmic proteins (FEA1/2) (100). We found a strong reduction of components of both Fe import systems in thiamine-induced *fdx1-rs* on NO_3_^−^ (Figure S9). The genes coding for the Fe(II) import proteins FRE1 and IRT1 were also found to be reduced on NH_4_^+^, but the Fe(III) import system and the FEA proteins were not affected on this N source. Lastly, IRT1 was the only protein involved in Fe assimilation whose expression was already reduced in *fdx1-1* knockdowns, while *FRE1* was also found to be reduced in *fdx2* mutants (Figure S8). No component of Fe import was affected in thiamine-uninduced *fdx1-rs*. We also analyzed the Fe content of different FDX mutants and knockdowns directly and found that Fe was significantly, albeit not dramatically, reduced only in thiamine-induced *fdx1-rs*, mirroring the results of the transcripts (Figure S9).

Taken together, these findings indicated to us that Fd reduction alone is not sufficient to cause a change in the Fe quota, but Fe assimilation is progressively reduced with increasing FDX1 depletion, first targeting the Fe(II) import system. At very low Fd content both pathways are diminished, but at this point of FDX1 reduction we already noted reduced expression of other highly abundant Fe proteins (especially PSI), which substantially contribute to the Fe quota. Despite reduction of cellular Fe there appears to be a deficiency in S supply towards Fe-S cluster biogenesis with severely FDX1 levels. To avoid detrimental effects of free Fe accumulating in cells, Fe import may need to be reduced. This may be a downstream effect of a lack of S assimilation, which may be at the root of the issue in *fdx1* knockdowns, including Fe-S cluster assembly.

### Other Fd dependent pathways are largely unaffected in *fdx1* and *fdx2* mutants

Fds are involved in cyclic electron transport, electron delivery from PSI to FNR for NADPH production and contribute to Chl biosynthesis at two steps, the synthesis of protochorophyllide from Mg-protoporphyrin IX ME (*CTH1*/*CRD1*) and Chl a to b interconversion via chlorophyll *a* oxygenase (*CAO*). FNR expression was not significantly affected in any mutant or condition (Data S3). Genes involved in tetrapyrrole biosynthesis and photosynthetic light and energy capture were mostly unchanged in their expression in thiamine-uninduced *fdx1-rs* and *fdx1-1*, with a few notable exceptions (Figure S5C). Expression of *CHLM* and *POR* were reduced in expression in different mutants, both on NO_3_^−^/NH_4_^+^, and two LHCII antenna proteins (*LHCBM4/8*). *CHLM* produces the Mg-protoporphyrin IX ME for the cyclase, while *POR* uses the product of the cyclase reaction, protochlorophyllide, as a substrate. CRD1, the lesser of two expressed cyclase genes predominantly used in Cu deficiency, was found reduced in abundance in thiamine-induced *fdx1-rs*, and was trending towards reduction in the other mutants. Chl content, and Chl a/b ratio were only affected in thiamine-induced *fdx1-rs/fdx2* double mutants (Figure S5B), with wildtype levels of Chl in other strains and conditions. In thiamine-induced *fdx1-rs* we noted a significant reduction for most proteins involved PSII/PSI and their antennas, while nuclear-encoded components of the Cyt b_6_f complex and chloroplast ATP synthase were not significantly changing but also trended towards reduction (Figure S5). Enzymes contributing to the CBB cycle were not significantly reduced, but similar to Cyt b_6_f complex and chloroplast ATP synthase were mildly reduced in induced *fdx1-rs* (Figure S5A, Data S3).

Taken together, expression of photosynthetic genes was mostly unchanged up until an almost complete removal of FDX1 from the chloroplasts. In induced *fdx1-rs*, most genes encoding for photosynthetic complexes are consistently reduced.

### DISCUSSION

FDX1/PetF and FDX2 are highly homologous proteins, sharing about 80% identical amino acids. The two Fds differ in the number of charges, surface charge distribution and midpoint potential (more negative for FDX1), which affect the kinetics of supported enzymatic reactions (Figure 1A) (59, 86). In addition to distinct transcriptional regulation (59)(Figure 3B) and their stoichiometry (Figure 4C), affinity towards potential Fd clients is largely reliant on ionic interactions (86, 101, 102), making these differences meaningful for each Fds *in vivo* client portfolio.

### FDX2 does not contribute to NADPH production

FDX1, the predominant chloroplast Fd isoform in Chlamydomonas, represents the bulk of the total Fd pool and is consistently responsible for NADPH generation through electron transfer to FNR (24). While *in vitro* enzyme kinetic studies indicated a higher affinity (lower K_m_) for FDX2 compared to FDX1 towards FNR, with similar turnover rates (k_cat_), FDX1 produced more NADPH by outcompeting FDX2 for binding to FNR, even when FDX2 is provided in excess of FDX1, likely driven by their difference in surface charges (24, 59, 86). The same study also found that FDX2 exhibits a higher affinity for soluble FNR than for thylakoid/PS1-associated FNR (24), indicating a preference of FDX2 for FNR for receiving electrons. *In vivo*, in photoheterotrophic conditions, FDX1 depletion did not significantly reduce the expression of FNR, or GAPDH, the main chloroplast NADPH client in the CBB, even when FDX1 was fully removed in thiamine-induced *fdx1-rs* (Figure S5, Data S3). Instead, expression of the photosynthetic complexes was reduced to ∼50 % in fully induced *fdx1-rs,* while it was not altered when there was still some residual FDX1 present (Figure S5A). This avoids excess electron delivery to a much smaller Fd pool in *fdx1* null mutants but leaves in place FNR for other aspects of metabolism, potentially for reduction of other Fds from NADPH. 6-phosphogluconate dehydrogenase (*GND1*) or Glucose-6-phosphate-1-dehydrogenase (*GLD1/2*) in the pentose phosphate pathway, or the glycolytic GAPDH (*GAPC1*), could provide NADPH to FNR in the Chlamydomonas chloroplast. While their expression is not significantly increased in any mutant analyzed here, we did note a minor increase of the pentose phosphate pathway enzymes in *fdx1-1* and thiamine-induced *fdx1-rs* mutants, but no change in *fdx2* (Figure S5). In the chloroplast of NO_3_^−^ grown Chlamydomonas cells FDX2 accumulated to ∼10% of FDX1 levels (Figure 2C), an abundance ratio that likely further drives preferential interaction between FNR and FDX1 *in vivo* (24). In this context it is also interesting to note that phototrophically grown *fdx2* mutants exhibit no growth defect when grown in either nitrogen source (Figure 3D). Transcriptome changes are very limited in *fdx2* mutants in general, again not affecting FNR, photosynthesis or the CBB.

Land plants like Arabidopsis have evolved two major, photosynthetic Fd isoforms, Fd1 and Fd2, that distribute electrons to FNR. A loss-of-function mutant in the predominantly expressed isoform, AtFd2, shows growth retardation but the plants grow and fix CO_2_ surprisingly well using the other isoform, Fd1, which represents around 5% of Fd levels in wildtype leaves (103). With FDX2 not involved in NADPH production on NH_4_^+^, this should make *fdx1* mutants more susceptible to growth conditions that don’t provide any reduced C (phototrophy). Chlamydomonas is a facultative phototroph that has been used extensively for the study of photosynthesis, utilizing mutant screens to obtain acetate-requiring mutants, defective in proteins required for photosynthesis (104–107). *fdx1* mutants are not acetate-requiring, they are not rescued by acetate addition. Furthermore, photosynthetic mutants are often light sensitive, requiring growth in low light conditions to produce biomass (106, 108). All different *fdx1* knockdowns analyzed here were grown in ambient light settings (∼ between 70 and 200 PAR), and while there are phenotypes with respect to growth rate, cultures with < 1% FDX1 abundance reached stationary culture densities at those light intensities, not too far behind wildtype growth rates. Even more surprising however, the phenotype of *fdx1* mutants was equally pronounced, if not more substantial, in photoheterotrophically-grown *fdx1* mutants, which are non-viable despite the availability of a reduced carbon source and less reliance on photoreduction for carbon assimilation (Figure 4D, S4C/E). This indicates that the growth defect in *fdx1* mutants is not from lack of supply to FNR for CO_2_ assimilation, but from other Fd-dependent reactions. Notably, *fdx1* mutants that express FDX2 (NO_3_^−^) are also less symptomatic. This indicates that partially overlapping functions between FDX1/2 are not with regards to electron distribution towards FNR and CO_2_ fixation, but to a different aspect of Fd-dependent metabolism that is not exacerbated by requiring photosynthetic growth.

Taken together, the data collected here further substantiates earlier *in vitro* studies at the organismal level, indicating that *in vivo* FDX2 is not involved in distributing electrons from PSI to FNR for NADPH production.

### Disparate roles for the major Fd FDX1 in N assimilation

FDX2 association with soluble FNR is indicative for receiving electrons from FNR, like in root-type Fds of land plants, where the main function of these specialized Fds is in NO_3_^−^ assimilation. While FDX2 is less abundant than FDX1 and *in vitro* has a similar affinity (K_m_) for nitrite reduction (59), we only found FDX2 in CoIP precipitates from NiR-HA tagged lines. Fd interactions with clients are transient, especially for nitrite reduction where 6 electrons are needed to be delivered sequentially. While these interactions might be hard to capture in cell lysates, capturing FDX2 but not FDX1 indicates a preferential recognition of FDX2 by NiR when both FDX1/2 are available in chloroplasts for electron supply. This is in line with the earlier observation that FDX2 supports nitrite reduction more efficiently *in vitro*, and changes in amino acid composition in FDX2 that revert differences with FDX1 affecting catalytic properties negatively, mostly affinity (K_m_) (86). In a crowded stroma *in vivo*, where both Fd’s are present, increased catalytic rates already appear selective to ensure FDX2 usage over FDX1 at NiR, even at an abundance disadvantage. While FDX2 is used for nitrite reduction, it is not required for growth on nitrate, FDX1 can cover that aspect of Fd-dependent metabolism when necessary (Figure 5C). Its also interesting to note that no additional FDX1 is required to compensate for FDX2 function, FDX1 appears to be present in excess in Chlamydomonas cells, which is also indicated by a large amount of reduction of FDX1 (> 95% reduction in various knockdowns) still producing viable cultures. Nitrate-grown cultures are even more resilient to FDX1 knockdown.

We therefore propose that FDX2’s contribution to Chlamydomonas Fd-dependent metabolism is to allow FDX1 to maintain its client distribution structure independent of the available N source. While other Fds are available in any mutant or knock-down strain analyzed here, all other chloroplast Fd’s in *fdx1* or *fdx2* mutants are either expressed at wild-type levels or less (Figure 6C). Presence of other Fds does also not result in any growth on nitrate in *fdx1/2* double mutants (Figure 5C). While there is some functional overlap between FDX1 and 2 in N assimilation, these quite similar Fds appear to control their set of Fd-dependent chloroplast metabolism, without interference from other Fds. This is further shown by the absence of effects on redox regulation, fatty acid synthesis and, mostly, Chl biosynthesis, which is affected in FDX1 knockouts, but not prior to full depletion.

To summarize, while FDX2 seems not to contribute to NADPH production, it is the preferred electron donor to NiR. But FDX1 is able to distribute electrons to NiR in its absence.

### S assimilation cannot keep up with intracellular demand

When FDX1 is fully removed, Chlamydomonas cells eventually cede to grow, most interestingly on NH_4_^+^ (no FDX2 expression, no nitrite reduction required) and in the presence of the reduced carbon source acetate. We opted for an analysis of various levels of FDX1 reduction in different knockdowns to identify Fd-dependent pathways already affected at sub-lethal levels. CO_2_ or N assimilation are likely not responsible for phenotypes in strains with some residual FDX1, as photosynthetic gene expression is only affected by a full removal of FDX1. Nitrite reduction is either not required (especially when NH_4_^+^ is provided) or preferentially handled by FDX2. When surveying other Fd-dependent metabolism, Fd-dependent sulfite reduction became the likely candidate to cause the growth defect in *fdx1*. This hypothesis is supported by a reduced S content in *fdx1* mutants, which is FDX2-dependently partially restored (Figure 7C). In addition, *fdx1* mutants induce expression of genes involved in increasing S affinity, especially importers activated in S deficiency, and aspects of S sparing, for example the remodeling of the cell wall with low S containing cell wall proteins (Figure 7AB). The composition of transcriptional targets mirrors the response that has been previously observed in S starved Chlamydomonas cells (72), despite luxurious amounts of S being available to *fdx1* strains. We got the impression that, while the number of differentially expressed genes in S-starved and FDX1-starved cells is similar, the amplitude of induction observed in S limited cells exceeded the one observed in *fdx1* knockdowns.

In plants, SiRs catalyze the reduction of sulfite to sulfide in chloroplasts using Fd as an electron donor. However, while plants like Arabidopsis and maize contain a single, Fd dependent SiR (4, 109, 110), Chlamydomonas harbors three genes coding for proteins involved in sulfite reduction (*SiR1, SiR2* and *SiR3*). While none of the sulfite reductases have been studied in detail in Chlamydomonas, SiR1 and SiR2 are highly similar to the monomeric, plant-type, Fd-dependent SiR from Arabidopsis, and to a lesser degree to the siroheme-containing hemoprotein of the multimeric NADPH-dependent sulfite reductase from *E. coli* (Figure S1C) (111–114). SiR3 is more similar to the diflavin reductase component of the multimeric *E. coli* complex (SiRFP), which receives two electrons at a time from NADPH, before channeling them to the catalytic subunits (SiRHP) of the complex (115). The presence of SiR3 indicates that Chlamydomonas may use a monomeric Fd-dependent SiR and a multimeric NADPH-dependent enzyme, which would require one or both of SiR1/2 to provide the catalytic domains to the complex. Observed transcriptional responses and growth defects in *fdx1* mutants could therefore be either the direct result of less efficient electron delivery to a Fd-dependent SiR by FDX1 or of reduced, FDX1-dependent NADPH production. We prefer the first scenario, since both SiR1/2 are increasingly expressed in response to FDX1 depletion, while the mRNA abundance of the NADPH-accepting SiR3 is reduced in *fdx1* (Figure 7A/B). The electron donor in either case would likely be FDX1, since S assimilation is independent of the N source, and mostly affected in *fdx1* mutants that don’t express FDX2 (NH_4_^+^). To make matters more complex, nitrite reductase and sulfite reductase are distantly related and are capable to utilize either nitrite or sulfite as a substrate *in vitro*, albeit with worse efficiencies for their respecitive secondary substrate (73, 74). FDX2-dependent rescue could therefore happen either by electron supply to SiR directly, or indirectly by sulfite reduction via NiR with FDX2 as the electron donor. In this case, both models seem equally likely based on our data, since S assimilation is still affected in nitrate grown *fdx1* null alleles in which FDX2 accumulates to ∼10% of FDX1 wildtype levels. This reduction in efficiency could either come from the lower affinity of NiR towards sulfite or FDX2 towards SiR. Interestingly, NiR was found to be de-repressed in response to S deficiency (72), while other components of nitrate/nitrite assimilation (nitrate reductase, transporters) were not. Induction of NiR in the presence of ammonium but absence of sulfur was magnitudes lower than on nitrate media, which would imply that sulfite reduction via NiR may be possible but is not a favored strategy.

FDX1 and FDX2 both can contribute to S assimilation, but there is a clear preference towards FDX1 as electron donor for this aspect of nutrient assimilation. Similar to nitrite reduction, no other Fd appears to be involved in S assimilation.

## ACKNOWLEDGMENTS

We thank the staff at the MSU Genomics Core for help with RNA quality control, cDNA sample preparation, library preparation and sequencing. We are thankful to the MSU proteomics core facility and Douglas Whitten for mass-spectrometry based protein identification of immunoprecipitated samples. We thank Sabeeha Merchant for kindly contributing Fd-specific antisera. This research was partially supported by a fellowship from Michigan State University under the Training Program in Plant Biotechnology for Health and Sustainability (T32-GM152798 and T32-GM110523). This work was also supported by grants from the National Institutes of Health for the resource entitled “Quantitative Elemental Mapping for the Life Sciences” (P41 GM135018) and from the U.S. Department of Energy, Office of Science, Basic Energy Sciences (DE-FG02-91ER20021). This work was supported in part through computational resources and services provided by the Institute for Cyber-Enabled Research (ICER) at Michigan State University.

## AUTHOR CONTRIBUTIONS

St.S. and D.S. designed the research; St.S., G.K.A., Sa.S., D.V. performed research; St.S., T.O. and D.S. analyzed and interpreted data; St.S. and D.S. wrote the paper. All authors edited and approved the final version of the manuscript.

## MATERIAL AND METHODS

### Generation of transgenic strains using LbCpf1/CRISPR gene editing

Arginine auxotrophic Chlamydomonas wildtype strain CC-425 was used to generate CRISPR/ Cpf1 gene edited strains as described in (Pham et. al 2022, Strenkert et. al 2024). In brief, cultures were grown to cell densities between 2 and 4 x 10^6^ cells/mL, assessed using a Beckman CoulterCounter Z2. Exactly 2 x 10^7^ cells were collected (3 min, 1424 x g) and resuspended in MAX Efficiency Transformation Reagent (Invitrogen, 1 mL), followed by suspension in 230 μL of the same reagent supplemented with sucrose (40 mM). Cells were incubated in a water bath at 40°C for 30 minutes. Purified LbCpf1 (80 μM) was preincubated with 2 nmol of the respective gRNA (gRNA target sequences are listed in Table 1) at 25 °C for 30 min to form RNP complexes. After incubation, 230 μL heat shocked cells were mixed with preincubated RNPs and linearized plasmid containing the respective selection marker. *ARG7* in pHR11, conferring the ability to grow without arginine supplementation was used in most cases, pSL18 containing the APHVIIIV gene referring resistance against paromomycin was used for the generation of *fdx1-rs fdx2* double mutants. To achieve template DNA-mediated editing, 4 nmol ssODNs were added (all ssODN templates are listed in Table 1). Cells were electroporated in a 4-mm gap cuvette (Biorad) at 600 V, 50 μF, 200 Ω by using Gene Pulser Xcell (Bio-Rad). Cells were recovered overnight in darkness and without shaking in 5 mL TAP with 40 mM sucrose and polyethylene glycol 8000 1% (w/v) and then plated after a 3 min centrifugation at RT and 1424 x g using the starch embedding method, using 60% corn starch.

Depending on the target, we used different methods to screen transformants for successful gene editing. To screen *fdx2* single and *fdx1-rs fdx2* double mutants harboring the two in-frame stop codons, we used colony qPCR. Colony qPCR of transformants was performed as follows: after one week of growth, half of each colony on TAP plates was resuspended in 100 uL 10 mM Tris-HCl (pH 8) 1 mM EDTA buffer in a 96 well plate. Cells were heated to 96 ***°***C for 10 min. After vortexing, 96 well plates were spun down for 4 min at ∼1000 x g. Supernatants containing the DNA were transferred to new 96 well plates. qPCR was performed using the oligos listed in Table 1. The oligos were designed to only result in either successful or unsuccessful PCR amplification if genome editing occurred, depending on the target. The following program was used for all qPCR reactions: 95°C for 5 min followed by 40 cycles of 95°C for 15 s, target specific annealing temperature for 30 s (between 58°C and 65°C) and 72°C for 30 s. Strains in which insertion of the *THI4* riboswitch was attempted were first screened on viability on agar plates. Colonies were transferred to new TAP plates either with or without 10 µM thiamine supplementation. Colonies showing reduced growth on either plates were used for further characterization.

Candidate transformants in which gene editing resulted in expression of a HA-fusion protein (FDX1-HA, FDX2-HA and NiR-HA) were screened directly at the protein level. Cells were grown in 1ml TAP medium with either ammonium (for FDX1-HA) or nitrate (FDX2-HA, NiR-HA) as the sole nitrogen source in 1.5 ml test tubes. Cells were collected (30 sec at 5000 x g, 4°C), the supernatant was discarded and the pellets were directly resuspended in 200 µl 2 x SDS sample buffer. Proteins were denatured for 20 min at 65°C and separated using SDS-PAGE, followed by transfer to nitrocellulose membranes and immunodetection using an HA specific antibody. Strains whose lysates showed a band from HA antibodies at the expected size were selected for further characterization.

All identified strains were further analyzed, first by Sanger sequencing, to confirm intended edits. For this purpose, transformants grown on TAP plates were used for another colony PCR using oligos amplifying 200-400 bps around the gene editing site (Table 1, sequencing oligos). Because we had difficulties amplifying the gene editing site in *fdx1-1* and *fdx1-rs* mutants due to their relatively large insertions, we isolated genomic DNA for whole genome sequencing instead. Sequences were mapped back to the Chlamydomonas genome (v6.1) either in its native assembly or with expected edits. For *fdx1-1* and *fdx1-rs* a break in coverage was found in the 5’UTR of the *FDX1* gene close to the used PAM site. ssODN templates were inserted into these breaks (*fdx1-rs*) and reads were mapped back to the modified genome confirming successful integration of the THI4 riboswitch. For *fdx1-1*, reads partially mapping to the insertion site were retrieved to recover the inserted sequences, then the genome was modified to account for the inserted sequences, before reads were mapped back to the locus. This process was repeated until the complete coverage was achieved.

### Culture conditions

Unless stated otherwise, cultures were grown either photoheterotrophically in Tris-acetate-phosphate (TAP) growth medium, or phototrophically in Sueka High Salt medium (HSM) supplemented with high (5%) CO_2_, in an Infors incubator with constant agitation (120 rpm) at 24°C in continuous light, photoheterotrophically at 65 µmol m^-2^ s^-1^ or phototrophically at 130 µmol m^-2^ s^-1^, provided by white LEDs. Both, TAP and HSM growth medium was prepared using the revised trace element formula (Special K) instead of Hunter’s trace element mixture (Kropat et al., 2011).

### Protein analyses by immunodetection

For analyses of total proteins, 2×10^7^ Chlamydomonas cells (at culture cell densities between 1-5 x 10^6^ cells/ml) were collected by centrifugation at 2500 x g at 4°C. The resulting cell pellet was resuspended in 50 µl of a buffer composed of 10 mM Na-phosphate buffer (pH 7) containing EDTA-free Protease inhibitor (Pierce, A32955). Samples were stored at -80°C prior to analysis. Protein amounts were determined using Pierce BCA Protein Assay Kit against BSA as a standard and diluted to 1-2 µg/µL with 2 x SDS sample buffer (125 mM Tris-HCl pH 6.8, 20% Glycerol, 4% SDS, 10% β-Mercaptoethanol, 0.005% Bromphenolblue). Proteins were denatured for 20 min at 65°C, separated on SDS-containing polyacrylamide gels using 20 μg of protein for each lane, unless stated otherwise. The separated proteins were then transferred by semi-dry electro-blotting to nitrocellulose membranes (Amersham Protran 0.45 NC). The membrane was blocked for 30 min with 3% dried non-fat milk in phosphate buffered saline (PBS) solution containing 0.1% (w/v) Tween 20. The same buffer was used as the diluent for both primary and secondary antibodies, which were incubated for at least 1h each. The membranes were washed in PBS containing 0.1% (w/v) Tween 20. Antibodies directed against CF1 (1:100 000), a polyclonal serum generated against native FDX1 purified from *Spinacia oleracea* (AS06 121, 1:1000), polyclonal serum originally generated by Terauchi *et al.* 2009, against the C-terminus of Chlamydomonas FDX1 (available through Agrisera AS11 1757, 1:5000) or FDX2 (AS16 3114, 1:1000), the HA epitope (Sigma, H6908, 1:15000) were used for immunodecoration. As secondary antibody, used at 1:15.000, we used Goat anti-Rabbit IgG, conjugated to alkaline phosphatase (Invitrogen, 31340) and detection was performed according to the manufacturer’s instructions.

### Co-Immunoprecipitations

Approximately 1×10^9^ Chlamydomonas cells (at mid-late log culture cell densities between 4-8 x 10^6^ cells/ml) were collected by centrifugation at 2500 g for 3 min at 4°C. Cell pellets were washed twice, first with 50 ml KH buffer (20 mM Hepes-KOH pH 7.2 and 80 mM KCl) in 50 ml conical tubes before being transferred to 15 ml conical tubes and washed with 15 ml of KH buffer. The resulting cell pellet was resuspended in 1 mL of lysis buffer with reduced ionic strength (20 mM Hepes-KOH pH 7.2, 1 mM MgC_l2_, 1 mM KCl, 15 mM NaCl, 1x protease inhibitor cocktail (Pierce, A32955), and 0.1% triton X-100) with 2 mM DSP (dithiobis (succinimidyl propionate), Lomant’s reagent). The cells were then broken by sonication on ice (25 x 1s intervals at 50% intensity, interspersed with 1s breaks). Cell debris was removed by centrifugation at 2500 g for 3 min minutes at 4°C, a 50 µl fraction of the cleared material was kept for input control purposes. The supernatant was transferred to a 1.5 mL centrifuge tube and combined with Protein A Sepharose beads coupled with anti-HA antibodies (Sigma, H6908). The protein-bead mixture was incubated with constant rotation at 4°C for 1 hour. The beads were then recovered by centrifugation at 5000 rpm for 20 s at 4°C, washed 4 times with lysis buffer, followed by two washes with 10 mM Tris-HCl (pH 7.6), each time using 5000 rpm for 20 s at 4°C to collect the cell pellet prior to resuspending the pellet in 1 ml of the indicated washing solution. The beads were transferred to a new 1.5 mL centrifuge tube in lysis buffer just prior to the switch in washing solution from lysis buffer to 10 mM Tris-HCl, to avoid carrying over unspecific precipitates not bound to beads. The last washing step in 10 mM Tris-HCl was used to split the precipitate into two equal sized aliquots. One aliquot was collected again and 50 µl 2 x SDS sample buffer were added to the ∼equal volume of wet beads for subsequent analysis by SDS-PAGE. The supernatant of the other aliquot was completely removed using a narrow, gel loading tip and the dry beads were frozen for subsequent protein identification by mass spectrometry (MS/MS). For proteolytic digestion, antibody-bound proteins were digested on-bead after washing 3 times using 50 mM ammonium bicarbonate. Trypsin, in the same buffer, was then added to the beads at 5 ng/µL so that the beads were just submerged in digestion buffer and allowed to incubate at 37°C for 6h. The solution was acidified to 1% with trifluoroacetic acid and centrifuged at 14,000 g. Peptide supernatant was removed and concentrated by solid phase extraction using StageTips (116). Purified peptides eluates were dried by vacuum centrifugation and frozen at -20°C or re-suspended in 2% acetonitrile/0.1%TFA to 20 µL. For LC/ MS/MS analysis, an injection of 10 µL was automatically made using a Thermo (www.thermo.com) EASYnLC 1200 onto a Thermo Acclaim PepMap RSLC 0.1mm x 20mm C_18_ trapping column and washed for ∼5 min with buffer A (99.9% Water / 0.1% Formic Acid). Bound peptides were then eluted over 35 min onto a Thermo Acclaim PepMap RSLC 0.075 mm x 250 mm resolving column at a constant flow rate of 300 nl/min with a gradient of 8% buffer B (80% Acetonitrile / 0.1% Formic Acid / 19.9% Water) to 22% B from 0 min to 19 min and 22% B to 40% B from 19 min to 24 min. After the gradient the column was washed with 90% B for the duration of the run. Column temperature was maintained at a constant temperature of 50°C using an integrated column oven (PRSO-V2, Sonation GmbH, Biberach, Germany). Eluted peptides were sprayed into a Thermo-Scientific Q-Exactive HF-X mass spectrometer (www.thermo.com) using a FlexSpray spray ion source. Survey scans were taken in the Orbi trap (60000 resolution, determined at m/z 200) and the top 15 ions in each survey scan were then subjected to automatic higher energy collision induced dissociation (HCD) with fragment spectra acquired at a resolution of 7500. The resulting MS/MS spectra were converted to peak lists using Mascot Distiller, v2.8.5 (www.matrixscience.com) and searched against a protein sequence database containing the Chlamydomonas genome (v6.1) appended with common laboratory contaminants (downloaded from www.thegpm.org, cRAP project) using the Mascot searching algorithm, v 2.8.3. (117). The Mascot output was then further analyzed using Scaffold, v5.3.3 (www.proteomesoftware.com) to probabilistically validate protein identifications. Assignments validated using the Scaffold 1% FDR confidence filter are considered true. Mascot parameters for all databases were as follows: allow up to 2 missed tryptic sites, variable modification of Oxidation of Methionine, peptide tolerance of +/- 10ppm, MS/ MS tolerance of 0.02 Da, FDR calculated using randomized database search.

### RNA extraction

3 x 10^7^ cells (at culture cell densities between 1-5 x 10^6^ cells/ml) were collected by centrifugation for 3 min at 2500 x g, 4°C. Resulting cell pellets were directly resuspended on ice in 1 ml of Trizol reagent, before being mixed with 200 µl chloroform : isoamyl alcohol (24:1). After rigorous vortexing, the samples were centrifuged for 15 min at 15 000 x g, 4°C. The supernatant was collected and total RNA was precipitated in cold ∼50% isopropanol. All RNA samples were subsequently treated with DNase I and concentrated / cleaned with the Zymo Research RNA Clean & Concentrator™-5 Kit according to the manufacturer’s instructions.

### Quantitative real-time PCR

Reverse transcription was primed with oligo dT(18) using 2.5 µg of total RNA and SuperScript III Reverse Transcriptase (Invitrogen) according to the manufacturer’s instructions. The resulting cDNA was diluted 15-fold before use. qRT-PCR reactions contained 5 μL of cDNA corresponding to ∼42 ng of total RNA, 10 pmol of each forward and reverse oligonucleotide, and 10 μL of iTaq Universal SYBR Green Supermix (Bio-Rad 1725122) in a 20 μL volume. Primers were optimized for 65°C melting temperature and amplicon sizes between 100 and 300 nt using Primer3 (118, 119). Primers targeting *RACK1* (Cre06.g278222_4532, previously referred to as CBLP) were used as loading control. The following program was used for all qRT-PCR reactions: 95°C for 5 min followed by 40 cycles of 95°C for 15 s, and 65°C for 60 s. Fluorescence was measured at the end of each 65°C cycle. A melting curve analysis was performed at the conclusion of the cycles from 65 to 95°C with fluorescence reads every 0.5°C confirming production of a single product.

### RNA sequencing

cDNA libraries were prepared from DNaseI treated, cleaned and concentrated (Zymo Research RNA Clean & Concentrator™-5) RNA preparations using the KAPA HyperPrep mRNA Library Preparation Kit with KAPA UDI indexes following manufacturer’s recommendations. Completed libraries were QC’d and quantified using a combination of Biotium AccuGreen High Sensitivity dsDNA and Agilent 4200 TapeStation HS DNA1000 assays. The libraries were pooled in equimolar amounts and the pool quantified using the Invitrogen Collibri Quantification qPCR kit. The library pool was loaded onto one lane of an Illumina S4 flow cell and sequencing was performed in a 2×150bp paired end format using a NovaSeq v1.5, 300 cycle reagent kit. Base calling was done by Illumina Real Time Analysis (RTA) v3.4.4 and output of RTA was demultiplexed and converted to FastQ format with Illumina Bcl2fastq v2.20.0. Reads in FastQ files were aligned to the Chlamydomonas genome (v6.1) using STAR v2.7.11b (120). SAMtools v1.18 was used for file format changes and sorting (121, 122), duplicate reads were removed using Picard Tools v3.0.0. Cufflinks was used to summarize reads and obtain expression estimates, DESeq2 was used for differential expression analysis.

### Statistical Analyses

Unless stated otherwise, a one-way ANOVA was used to test for differences between samples. Successful ANOVA was followed by a Holm-Sidak Post-hoc Test. Asterisks indicate a p-value of <0.05.

